# Dietary resilience of coral reef fishes to habitat degradation

**DOI:** 10.1101/2024.05.22.595166

**Authors:** Friederike Clever, Richard F. Preziosi, Bryan Nguyen, Brígida De Gracia, Helio Quintero, W. Owen McMillan, Andrew H. Altieri, Aaron O’Dea, Nancy Knowlton, Matthieu Leray

## Abstract

The ability of consumers to adjust their diet in response to resource shifts is a key mechanism allowing the persistence of populations and underlying species’ adaptive capacity. Yet on coral reefs, one of the marine habitats most vulnerable to global change, the extent to which species alter their diet remains poorly understood. Here, we integrated DNA-based gut content analyses (metabarcoding), otolith analysis, body condition, and field surveys to test how diet can mediate effects of habitat degradation on two invertivorous fishes: *Chaetodon capistratus*, a browser, and *Hypoplectrus puella*, an active predator. Metabarcoding revealed significant dietary variation in both species across a habitat gradient. However, the response was more pronounced in the browser, whose diet was anthozoan-dominated on healthy reefs, whereas annelid-dominated on degraded reefs. We found reduced growth and body condition on degraded reefs in the browser but not the active predator. Our results reveal that dietary versatility can serve as a mechanism to cope with degraded environments, but that species differ as to whether these changes are sufficient to buffer from changes in habitat. We detected intraspecific dietary variation across sites that suggests food webs and energy flow differ at relatively small scales between healthy and degraded reefs.

## INTRODUCTION

The capacity to take advantage of alternative resources is increasingly recognized as an important trait for species’ resilience in the face of habitat degradation (MacNally, 1995; Wong & Candolin, 2015). Such changes in feeding behavior, in turn, influence community dynamics (e.g., mediated by relative levels of niche overlap among species) and trophic interactions which underpin ecosystem functioning. Despite implications for both population persistence and ecosystem functioning, little is known about dietary versatility in the wild in response to changing resource landscapes and the potential consequences for consumers. This is particularly true in marine environments, where empirical studies of species dietary niches are challenging due to the complexity of marine food webs and the difficulty of documenting trophic behaviors (Donelson et al., 2019). However, with evolving high-throughput sequencing technology it is now possible to analyze large quantities of samples simultaneously, illuminating diets at unprecedented taxonomic resolution and spatial scale (Alberdi et al., 2019; Pompanon et al., 2012).

On coral reefs, fishes are subjected to accelerating rates of severe habitat change and degradation. This has led to declines in abundance across most trophic groups (Pratchett et al., 2018). Specialized species, such as obligate corallivores, are thought to be the most vulnerable (Wilson et al., 2006; Graham, 2007). Generalized feeding strategies, as observed in some species of coral reef fishes, may be more resilient to changes in resource availability (Wilson et al., 2006), but to which degree trophic versatility provides population resilience to habitat degradation remains poorly understood. This is in part because alternative prey choice may entail lowered nutritional uptake and thus reduce fish health condition with potential negative consequences for fitness and population persistence (Pratchett et al., 2004; Berumen et al., 2005; Hempson et al., 2017). Describing complex consumer-resource interactions has been hampered by limited empirical data at adequate prey taxonomic resolution for detecting potentially subtle dietary variation (Parravicini et al., 2020), especially in generalist feeders with flexible diets. As a consequence, more detailed knowledge of dietary resource use is required to understand responses to habitat and prey community change in coral reef fishes.

Here we leverage dietary metabarcoding (i.e., the DNA-based characterization of prey communities in stomachs and guts) to assess levels of dietary versatility of two common coral reef fishes across a habitat gradient in a Caribbean system. Fish diet information derived from gut contents has been commonly studied visually by describing the morphological features and hard-part remains of prey organisms (Baker et al., 2014; Nielsen et al., 2017; Traugott et al., 2020), and/or with behavioral observations of foraging and bite rates (Baker et al., 2014; Hyslop, 1980; Pratchett, 2005). The main advantage of dietary metabarcoding over these conventional visual approaches is that it allows identification of semi-digested, soft bodied, small, and/or cryptic organisms (meio- and microbiota) (Chariton et al., 2015; Leray & Knowlton, 2015) that may be missed by visual methods (Berry et al., 2015; Nagelkerken et al., 2009). Increased taxonomic resolution as facilitated by DNA-metabarcoding has already revealed higher levels of resource partitioning among closely related species than previously assumed (Brandl, Casey and Meyer, 2020; Leray, Meyer and Mills, 2015; Leray et al., 2019) and, conversely, unexpected patterns of dietary overlap (Coker et al., 2022). By enabling prey identification to an unprecedented taxonomic resolution, DNA metabarcoding allows detection of intraspecific variations in diet that would otherwise go unnoticed, and this in turn can be linked to changes in consumer populations and prey availability to understand the responses to habitat degradation.

We explored the dietary patterns of two common reef fish species with distinct feeding strategies, the browsing butterflyfish *Chaetodon capistratus* and the active predator hamlet *Hypoplectrus puella*, across nine reefs that vary in coral cover and benthic composition in the Bay of Almirante, situated on the Caribbean coast of Panama. We quantified links between diet (composition and breadth), fish age, growth and body condition, and prey densities across a habitat gradient created by severe hypoxic events in the Bahía Almirante in Bocas del Toro, Panama, providing conditions of a natural experiment (Altieri et al., 2017; Leray et al., 2021). Based on the literature (Birkeland & Neudecker, 1981; Gore, 1984) and on feeding observations (F. Clever, unpublished data), we hypothesized that *C. capistratus* would switch from an anthozoan dominated diet on high coral cover reefs to a broader suite of prey taxa on reefs where coral cover is very low. In contrast, we expected *H. puella* to maintain a high proportion of crustaceans in its diet across all reefs (e.g., Whiteman, Côté, and Reynolds 2007), but with compositional variation as a function of change in benthic invertebrate assemblages in relation to coral cover.

## METHODS

### Study system

Bahia Almirante is a large (450 km^2^), semi-enclosed coastal lagoon in the Bocas del Toro Archipelago on the Caribbean coast of Panama (Fig. 1A). Its specific geomorphology, climate and human pressures contribute to occasional reductions in dissolved oxygen levels (see supplementary methods, section I). In 2010, a hypoxic stress event led to drastic coral cover decline and die-off (Altieri et al., 2017) that contributed to a gradient of habitat degradation as coral cover sharply decreased on reefs in areas inside the bay (Fig. 1A, ‘inner bay disturbed’ zone), but less so or not at all in other areas of the inner bay (Fig. 1A, ‘inner bay’ zone) and outside the bay (Fig. 1A, ‘outer bay’ zone; Fig. 1B). We tested how habitat affects diet and body condition of two species of reef associated, benthic-feeding fishes along this gradient (Figs. 1C, 1D, 1E and 1F). We selected three discrete reefs from each of three reef zones based on coral cover data: “outer bay” (high coral cover), “inner bay” (intermediate coral cover), and “inner bay disturbed” (very low coral cover) (n=9 reefs total, Figs. 1A and 1B). Other related factors, such as water quality and exposure likely covary with coral cover across the area and may contribute to relative prey availability (Collin et al., 2009; D’Croz et al., 2005; Kaufmann & Thompson, 2005).

**Figure 1.**
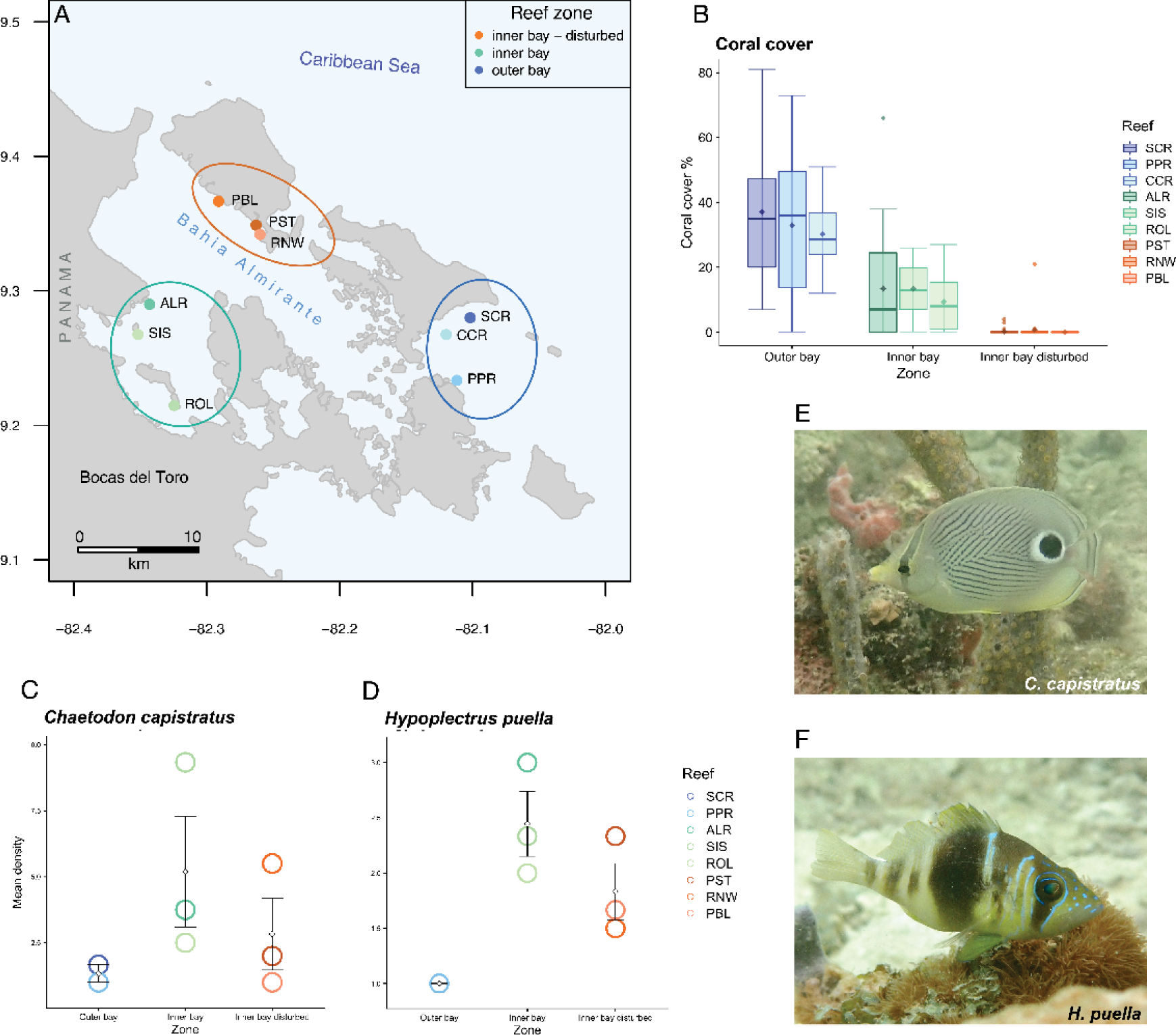
Study system in the Bahia Almirante, Bocas del Toro (Panama). (A) Map locating the nine study reefs across three zones characterized by different levels of live coral cover: outer bay reefs (blue) [Salt Creek (SCR), Cayo Corales (CCR) and Popa (PPR)]; inner bay reefs (green) [Almirante (ALR), Cayo Hermanas (SIS), and Cayo Roldan (ROL)]; inner bay disturbed reefs (orange) [Punta Puebla (PBL), Punta STRI (PST) and Runway (RNW)]. (B) Percent live hard coral cover across the habitat gradient from high coral cover (outer bay zone) to low coral cover (inner bay disturbed zone), boxplot upper and lower whiskers correspond to the first and third quartiles, bars depict medians, diamonds depict means. Fish density (mean ± sd) across reef zones for (C) *Chaetodon capistratus* and (D) *Hypoplectrus puella*. (E) *Chaetodon capistratus and (F) Hypoplectrus puella*. Photos: Matthieu Leray.

### Study species

Both study species are coral-associated, benthic carnivores that are common members of Caribbean reef fish assemblages (‘least concern’ conservation status, IUCN; Anderson et al., 2015; Rocha et al., 2010) (Figs. 1E and 1F). The barred hamlet *Hypoplectrus puella* (Cuvier) is a small sea bass (Perciformes: Serranidae) known to prey mainly upon crustaceans and to a lesser extent upon fishes (Holt et al., 2008; Randall, 1967; Whiteman et al., 2007) within small foraging territories, often in sit-and-wait mode (Barlow, 1975). The foureye butterflyfish *Chaetodon capistratus* (Linnaeus) feeds primarily by browsing on anthozoans with a preference for scleractinian corals (Birkeland & Neudecker, 1981; Liedke et al., 2018). It tends to feed almost continuously while visiting various colonies within a reef zone (e.g., crest), usually for short feeding bouts during which it nips on coral polyps (Pitts, 1991), but distances traveled may vary (Gore 1983).

### Benthic, fish and invertebrate surveys

To characterize habitat quality and fish populations, benthic cover and reef fishes were surveyed in May and June of 2016 at our nine study reefs. Coral cover levels remained stable between 2016 and our fish collections in 2018 (Doucette et al., 2022). Using three replicate 20 m transects per reef, we assessed benthic cover from photo quadrats, fish communities and study species’ densities along 20 x 5 m belts, and assessed diversity and abundance of macroinvertebrates (> 2 mm) within three quadrats per reef (50 x 50 cm) (sup. methods, section II).

### Fish collection

To characterize fish body condition and collect diet material, twenty adult fishes per species were collected at each of the nine reefs by spearfishing (Fig. 1A) in February and March of 2018 following protocols approved by the Institutional Animal Care and Use Committee of the Smithsonian Tropical Research Institute (IACUC). Immediately after spearing, each fish was anesthetized on the boat in a sterile and labeled Whirl-Pak bag with seawater and clove oil and subsequently stored on ice. Upon return to the field station, fish were weighted (g wet weight) and total length (mm TL) was measured using a digital caliper. Each fish was dissected under a laminar flow hood using sterile, DNA de-contaminated tools. Gastrointestinal tracts were individually preserved in 96% ethanol and stored at *−*20*^◦^C* until DNA extraction. Strict procedures were used to avoid cross-contamination (supplementary methods, section III).

### Otolith-based fish age determination and growth rate estimation

Pairs of sagittal otoliths were extracted from fish individuals (*C. capistratus* N = 158, *H. puella* N = 127) and photographed using a LEICA Model EZ4W stereoscopic microscope with an integrated camera and light system (Figs. S1A and S1B). One otolith of each pair was analyzed for annuli following the methods in Morales-Nin (1991) (Fig. S1C). Daily growth increments were examined for a subset of otoliths of each species to confirm that each annulus corresponds to one year of growth. The total length of each otolith (LO) was measured (*C. capistratus* N = 117, *H. puella* N = 127) while positioning the otoliths on their distal sides. We estimated fish growth rates by first assessing whether the relationship between otolith length and fish total length was linear, as would be expected. We then assigned the cut-off where the relationship between fish age and fish total length starts to become non-linear. Fish growth rates (mm year-1) were estimated for the linear segment of the fishes’ size range.

### Prey tissue preparation and DNA extraction

The digestive tract of each fish was separated into stomach and intestine. The stomach content represents a snapshot of the most recent prey ingested, whereas the intestinal content integrates semi-digested prey consumed up to multiple hours prior to collection. The stomachs of *C. capistratus*, and intestines of *H. puella*, were dissected longitudinally and contents and digesta isolated respectively. The stomachs in *C. capistratus* contained a diverse and representative assortment of prey items, presumably because of the species’ continuous browsing behavior. In contrast, the stomachs of the sit-and-wait occasional predator *H. puella* frequently contained only a single prey item, and we therefore sampled intestines that yielded more diverse representations of prey items. Prey tissue was removed from stomachs and digesta and mucosa isolated from intestines using sterile and DNA-decontaminated forceps and disposable sterile surgical blades. Gut mucosa was included here since samples were also used for analysis of bacterial communities (not presented in this study) (Clever et al., 2022). Isolated stomach and intestinal contents were then weighed (wet weight mg) individually on clean, sterile weighing boats on a digital scale. Dissection and extraction blanks were introduced at this step by performing each preparation step with a sample consisting of nuclease-free water. One negative control was included in each set of extractions (∼20 samples). DNA was extracted from between 0.05 and 0.25 g of prey tissue per sample using the Qiagen Powersoil DNA isolation kit following the manufacturer’s instructions with minor modifications to increase the yield (supplementary methods, section IV). DNA was eluted in 100 ul buffer (C6 solution).

### Metabarcoding library preparation

To enable identification of prey items to the species level, we targeted a 313 bp fragment of the hyper variable mitochondrial Cytochrome c Oxidase subunit I (mtCOI) gene region with a versatile PCR primer set (mlCOIintF and jgHCO2198; Geller et al., 2013; Leray et al., 2013a) (Table S1). This primer set, originally designed for the amplification of metazoan DNA, was shown to be effective at characterizing coral reef fish gut contents (Leray et al., 2013a), and has previously successfully amplified diverse bulk samples of marine benthic taxa as well as provided useful abundance estimates (Leray & Knowlton, 2015). In each PCR reaction, we included consumer-specific annealing blocking primers (Table S2) (at 10x COI primers), as amplification of consumer DNA can overwhelm the recovery of prey (Vestheim & Jarman, 2008). Blocking primer design and thermocycling parameters followed the methods described in Leray et al. (2013b). Polymerase Chain Reaction (PCR) was carried out for three replicates of each sample to enhance prey detection probability and account for variation in PCR amplifications caused by PCR drift (Alberdi et al., 2017; De Barba et al., 2014). We employed a PCR-free library preparation approach with matching tags using the TruSeq DNA PCR-free LT library Prep Kit (Illumina). Our methods for sample multiplexing, PCR reactions and library preparation are detailed in the supplementary methods (section V). Sequencing of the final product was performed on an Illumina MiSeq sequencer (reagent kit version 2, 500 cycles) at the George Washington University, Washington, DC.

### Sequence analysis

All analyses were conducted in R version 4.1.3 (R. Development Core Team, 2008). After demultiplexing, sequence reads were adapter-, primer- and quality-trimmed with Flexbar (version 3.0.3; Roehr, Dieterich & Reinert 2017). Subsequently, sequences were filtered, chimera-checked, and processed into amplicon sequence variants (ASVs) with DADA2 (Callahan et al., 2016). ASVs were then clustered with VSEARCH (Rognes et al., 2016) at a 97% identity threshold into OTUs to approximate biological species. OTUs were curated with the LULU algorithm (Frøslev et al., 2017) by reducing taxonomic redundancy and enhancing the richness estimate accuracy (LULU parameters: minimum ratio type = “min”, minimum ratio = 1, minimum match = 84, minimum relative co-occurrence = 0.95). OTUs were assigned taxonomy using the Bayesian Least Common Ancestor (BLCA) taxonomic classifier (Gao et al., 2017) against the Midori-Unique v20180221 database (Leray et al., 2022; Machida et al., 2017), which is a curated library of metazoan COI sequences (available at www.reference-midori.info). We omitted all BLCA taxonomy assignments of less than 50% confidence. Unassigned OTUs were identified using BLAST searches (word size = 7; max e-value = 5e-13) against the whole NCBI NT database (retrieved May 2018), and the lowest common ancestor of the top 100 hits was used to assign taxonomy. We retained all OTUs delineated as Metazoa for downstream statistical analysis. All OTUs delineated as either one of our fish study species were removed.

## Statistical analyses

### Benthic, fish and invertebrate surveys

Differences in mean percent coral cover among zones were assessed using Kruskal-Wallis test (Kruskal & Wallis, 1952) with Benjamini-Hochberg corrected post hoc Dunn’s Test (Benjamini & Hochberg, 1995; Dunn, 1964) (kruskal.test function, stats package v 4.1.3; dunnTest function, FSA package v 0.9.1; Ogle et al., 2020). For each reef, hard coral diversity (Shannon index) was estimated and visualized as boxplots. Differences in benthic composition among three zones were visualized using Principal Coordinates Analysis (PCoA) based on Bray-Curtis dissimilarity (Bray & Curtis, 1957). Eigenvectors were plotted depicting the relative contribution of benthic groups to separation among zones. Non-metric multidimensional scaling (NMDS, Clarke & Warwick, 2001) with Bray-Curtis dissimilarity was used to assess differences in fish communities among habitat zones. We visualized the densities of our fish study species across zones and tested for significant differences among zones using one-way Analysis of Variance (ANOVA; aov function, stats package v 4.1.3). Variation in mean densities of frequently consumed benthic prey taxa, selected based on metabarcoding results, were compared at various taxonomic levels (i.e., all sampled invertebrates, arthropods, decapods, true crabs [Brachyura], mithracid crabs, spaghetti worms [Terebellidae]) among zones using boxplots, and significant differences assessed by ANOVA with post-hoc Tukey test (aov and TukeyHSD functions, stats package v 4.1.3). Both overall invertebrate and arthropod community composition were visualized using stacked barcharts of relative densities.

### Fish length-weight and body condition

We first modeled the length-weight relationship by species for the whole dataset (linear regressions, lm function; FSA R package v 0.8.30; Ogle, Wheeler & Dinno 2020)

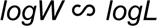

where L and W are the natural log transformed fish total length (mm) and weight (g), respectively. To compare fish body condition (e.g., relative ‘plumpness’ in relation to length—with plumper fish of a given length assumed to be in better condition; Tesch 1968; Froese, 2006) among zones, we calculated the relative condition factor for each individual (*Kn*) (Le Cren, 1951) by estimating the deviation between the observed weight to the predicted length-specific mean weight of the population (Blackwell et al., 2000; Froese, 2006)

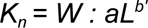

where a and b are the species- and population-specific length-weight parameters, respectively, obtained from the length-weight regression. L is the natural log transformed observed total length (mm) and W is the natural log transformed observed weight. Due to the local scope of our study focusing on small-scale spatial differences among fish subpopulations within species, the relative condition factor (Kn) was used as opposed to the relative weight (wr), the latter being based on standard weight developed across populations (Blackwell et al., 2000). To assess how fish condition varied across zones, we first tested if relative condition (*Kn*) varied among zones overall using Kruskal-Wallis tests. Second, we plotted relative fish condition *Kn* against three fish size classes (both species: 50 - 79 mm, 80 - 99 mm, 100 - 130 mm) grouped by zone and tested for significant differences within size classes among zones (Kruskal-Wallis test with post hoc Dunn’s Test). Lastly, we assessed whether the slopes of the length-weight regression differed among zones by modelling the interaction between fish total length and zone in affecting the length-weight relationship

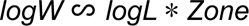

ANOVA was used to determine whether slopes differed significantly (anova function; FSA R package). In addition, we tested for significant differences in wet weight (g) and total length (mm) among zones (ANOVA; aov and TukeyHSD functions).

### Age and growth

We tested whether there were significant differences in fish age and growth rates among zones evident in otolith samples (Kruskal Wallis test with post hoc Dunn’s Test).

### Diet composition

NMDS ordination (Clarke & Warwick, 2001) based on Bray-Curtis dissimilarity was used to visualize differences in diet composition among reefs and zones for each fish species (isoMDS function, Mass package; Venables & Ripley, 2002). Stacked barcharts were generated for each fish depicting relative read abundances of prey OTUs across nine study reefs (phyloseq package; McMurdie & Holmes, 2013). We used generalized linear mixed effects models (GLMMs) with a negative binomial distribution (function glmer.nb, lme4 package v1.1-21; Bates et al., 2015) to test for effects of percent coral cover on sequence relative read abundance of dominant diet categories as identified by metabarcoding: (i) annelids and hard corals in *C. capistratus*, and (ii) benthic and planktonic crustaceans in *H. puella*. Final models were selected based on likelihood ratio tests against null models (supplementary methods, section VI). We then tested the effect of fish consumer age on the relative abundance of these main diet items. To characterize the feeding strategy of both fish species in terms of how specialized or generalized the diet appears on the population level, we used a graphical analysis proposed by Amundsen et al. (1996) modified from Costello (1990). To generate diagrams representing feeding strategy and prey importance at three reef zones, frequency of occurrence was expressed as percentage and calculated by dividing the number of fish individuals in which a prey item was present by the total number of fish. Prey specific abundance was calculated as the percentage of the diet that a food item represents across only those fish individuals where it was present

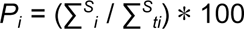

where *P_i_* is the prey-specific abundance of prey i, *S_i_* is the abundance of prey i in the stomach or intestines, and *S_ti_* is the total prey abundance in only consumer individuals where prey i is present. Because a generalist diet profile may arise at the population level from either broad individual diets and/or high variation in diet composition among specialized individuals (Amundsen et al., 1996; Bolnick et al., 2003), Amundsen’s method includes an indirect measure of the contribution to niche width of both within individual variation (within phenotype component, WPC) and variation among individuals (between phenotype component, BPC).

## RESULTS

### Benthic, fish, and invertebrate surveys

Percent live coral cover and coral diversity differed among the three reef zones. The outer bay zone featured the highest levels of live coral cover (percent cover per transect across three reefs mean ± SD = 33.46 ± 3.41, Fig. 1B) and coral diversity (Shannon diversity, Fig. S2). Live coral cover and coral diversity were lower at the inner bay zone (12.06 ± 2.3; Fig. 1B, Fig. S2), and live corals were nearly absent at reefs in the inner bay disturbed zone (0.34 ± 0.39; Fig. 1B), which also had the lowest levels of coral diversity (Fig. S2). Principal coordinates analysis (PCoA) of benthic composition revealed that differences among zones were driven by sponges and dead corals (inner bay disturbed zone), ‘other invertebrates’ (inner bay zone) and live hard and soft corals with macroalgae (outer bay zone) (Fig. S3). Mean fish density peaked at the inner bay zone for both species, but differed significantly only in the case of *H. puella* (*C. capistratus*: ANOVA; *F* = 1.8, *p* = 0.25 and *H. puella*: ANOVA; *F* = 1.8, *p* = 0.01) (Figs. 1C and 1D). NMDS ordination of fish communities showed clustering of reefs located at the outer bay zone and inner bay disturbed zone, respectively, whereas the inner bay zone was more variable (Fig. S4). Overall benthic invertebrate mean density significantly differed among three zones with highest levels at the outer bay zone (Fig. S5A, Table S3); post hoc testing confirmed a significant difference between the inner bay disturbed zone and the outer bay zone (Fig. S5A, Table S4). Regarding individual invertebrate groups constituting important fish prey, we found no significant differences in the mean densities of either spaghetti worms (family: Terebellidae, phylum: Annelida) (Fig. S5B, Table S3) or crustaceans (phylum: Arthropoda) (Fig. S5C, Table S3). Within Arthropoda, there were also no significant differences among zones for decapod crustaceans (order: Decapoda) (Fig. S5D, Table S3), Brachyura (true crabs, order: Decapoda) (Fig. S5E, Table S3), and mithracid crabs (family: Mithracidae) (Fig. S5F, Table S3). Crustaceans of class Malacostraca (phylum: Arthropoda) were more abundant at the inner bay disturbed zone than the outer bay zone (Fig. S6A). Within arthropods, the relative densities of decapod crustaceans (class: Malacostraca) were highest at the inner bay zone and at one site (PPR, Popa Reef) of the outer bay, but lower at reefs of the inner bay disturbed zone and Salt Creek Reef (SCR) at the outer bay (Fig. S6B).

### Fish length-weight relationship and body condition

Total length (TL) ranged from 53.49 to 98.19 mm (mean ± SD = 80.08 ± 10.73) for *C. capistratus,* and 56.73 to 125.23 mm (91.19 ± 8.96) for *H. puella;* wet weight (W) ranged from 5.34 to 34.40 gr (17.97 ± 7.25) for *C. capistratus*, and from 3.01 to 23.49 gr (14.60 ± 3.8) for *H. puella.* One-way ANOVA showed that fish length and weight differed significantly among zones for both species (ANOVA; *C. capistratus F* = 2889.54, *p* < 0.001; *H. puella F* = 1550.38, *p* < 0.001). *Chaetodon capistratus* were on average longer and heavier at the inner bay zone (TL mean ± SD = 87.00 ± 10.26; W mean ± SD = 23.04 ± 6.71) and shorter and lighter at the inner bay disturbed zone (TL = 73.02 ± 10.20; W = 13.41 ± 6.62), with intermediate values at the outer bay zone (TL = 80.33 ± 6.99; W = 17.59 ± 5.16). *Hypoplectrus puella* were on average longer and heavier at the inner bay disturbed zone (TL = 94.07 ± 6.56; W = 15.98 ± 3.08), whereas individuals were slightly shorter and lighter at the inner bay (TL = 92.03 ± 9.05; W = 14.55 ± 3.71) and outer bay (TL = 87.49 ± 9.78; W = 13.27 ± 4.35) zones. We found a significant interaction between fish total length and zone in affecting the length-weight relationship for *H. puella* but not *C. capistratus* (ANOVA; *C. capistratus F* = 2.04, *p* = 0.13; *H. puella F* = 9.12, *p* < 0.001). The relative fish condition factor (*Kn*) pooled across size classes did not differ among zones for either species (Kruskal-Wallis test: *C. capistratus*, *X^2^*= 2.41, *p* = 0.3; *H. puella*, *X^2^* = 2.36, *p* = 0.31) (Figs. S7A and S7B). However, when comparing *Kn* levels across zones within a given size class, we found that medium-sized *C. capistratus* differed among zones (*X^2^* = 8.41, *p* = 0.01, Fig. 2A). The other size classes showed no differences among zones (small: *X^2^* = 1.71, *p* = 0.43; large: *X^2^* = 4.56, *p* = 0.1, Fig. 2A). Post hoc tests confirmed that fish condition levels in the medium size class were significantly lower at the inner bay disturbed zone than at the inner bay (Dunn’s; adjusted *p* = 0.01), and lower in the outer bay than the inner bay (adjusted *p* = 0.05). The inner bay disturbed and outer bay zones did not significantly differ (adjusted *p* = 0.21) (Fig. 2A). *Hypoplectrus puella* showed no significant differences in body condition (*Kn*) within size classes among zones (small: *X^2^* = 3.36, *p* = 0.19; medium: *X^2^*= 3.66, *p* = 0.16; large: *X^2^* = 1.72, *p* = 0.42, Fig. 2B). *Chaetodon capistratus* showed a greater variability in condition within and among the three size classes at the inner bay disturbed zone compared to both other zones (Fig. 2A). For *H. puella* in the outer bay, condition in small fish appeared significantly lower than in large fish, whereas in the inner bay zone, condition was lower in large fish (Fig. 2B).

**Figure 2.**
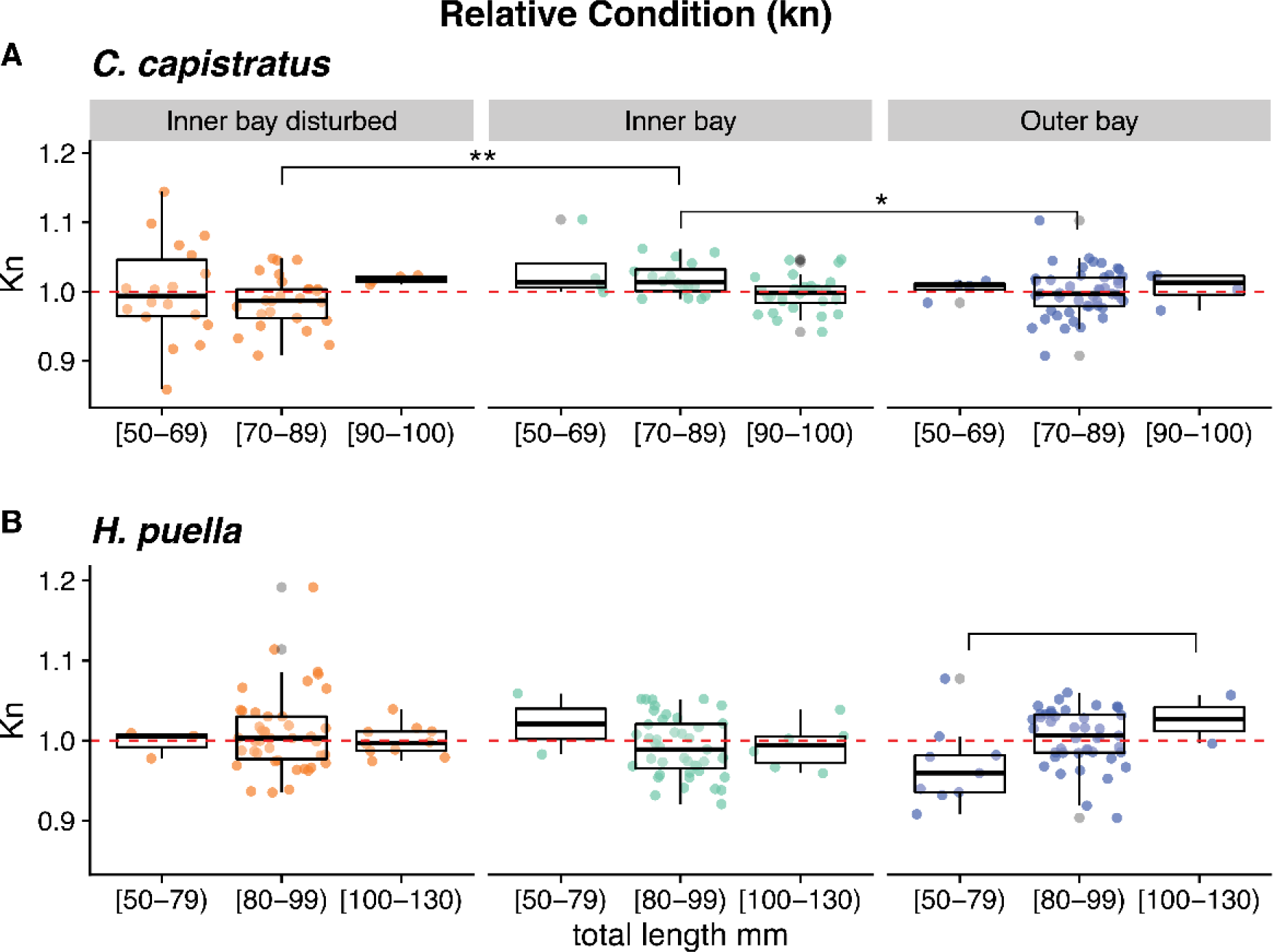
Relative Condition Factor (Kn) across three reef zones by fish size classes. Calculations are based on length-weight regression models (A) for *Chaetodon capistratus* and (B) for *Hypoplectrus puella*. Significant levels based on adjusted *p*-values are denoted as ** = 0.01 and * = 0.05. Significant differences among size classes within zones are based on comparing box plots and represented solely by brackets.

### Otolith-based fish age determination and growth rate estimation

Fish age ranged from three to nine years (mean ± SD = 5.8 ± 1.09) for *C. capistratus,* and from three to eight years (5.1 ± 0.94) for *H. puella.* On average, *Chaetodon* individuals were oldest in the inner bay zone (6.41 ± 1.42) in comparison to both the disturbed (5.46 ± 0.81) and outer bay (5.66 ± 0.84) zones with a significant difference in age among zones (Kruskal; *X^2^ =* 13.56*, p* = 0.001; inner bay vs inner bay disturbed zone: Dunn’s; adjusted *p* = 0.0009; inner bay vs outer bay zone: adjusted *p* = 0.01; inner bay disturbed vs outer bay zone: adjusted *p* = 0.28). *Hypoplectrus* individuals in our sample exhibited a similar age structure across the three zones: inner bay (mean ± SD = 5.1 ± 0.82), inner bay disturbed (5.07 ± 1.01) and outer bay (5.14 ± 1) showing no significant difference in age among zones (Kruskal, *X^2^ =* 0.12784*, p* = 0.94). Fish growth rates in individuals of age < 6 years differed significantly among three zones in *C. capistratus* (Kruskal, *X^2^* = 10.761, *p* = 0.005) but not *H. puella* (Kruskal, *X^2^* = 4.19, *p* = 0.13) (Fig. 3). *Chaetodon capistratus* individuals grew significantly slower at the inner bay disturbed than at the inner bay (Dunn’s; adjusted *p* = 0.007) and the outer bay (adjusted *p* = 0.03) zones.

**Figure 3.**
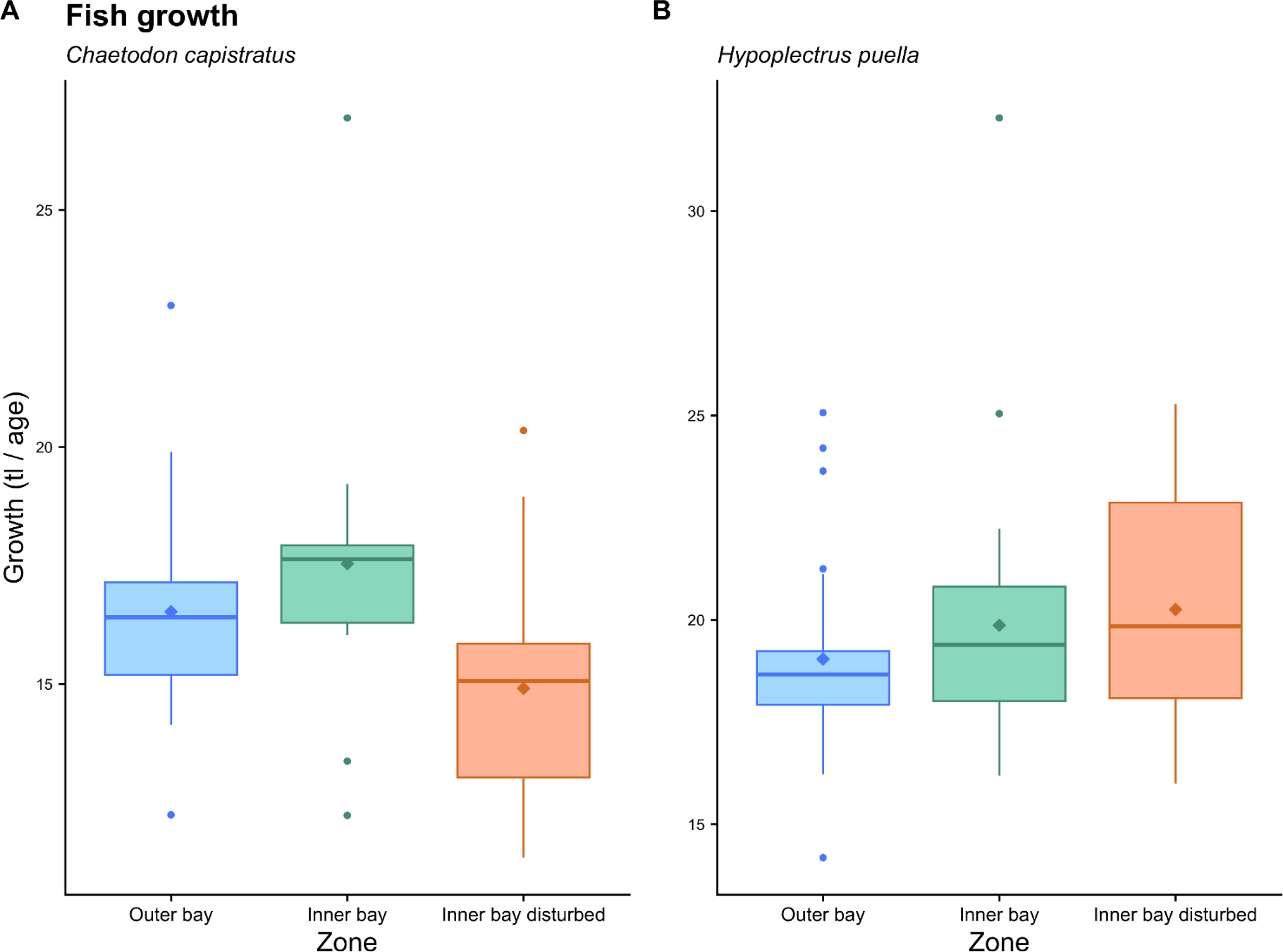
Differences in fish growth among three zones for (A) *C. capistratus* and (B) *H. puella*. Growth was estimated for fish individuals of ages <6 years using age data generated from our otolith analysis.

## Diet composition

### Sequence Analysis

A total of 18,427,824 raw paired-end reads were obtained. After denoising, removing chimeras, and processing, retained high quality reads clustered into 1009 OTUs assigned to the kingdom Metazoa, of which 166 (16.5%) were matched to species. An additional 613 OTUs (60.8%) could be assigned to higher taxonomic levels. The extraction and PCR controls did not show contamination. Steep sample-based rarefaction curves indicate that the two target species consume a large diversity of prey in each reef zone that exceeds the prey we identified in our sampling (Figs. S8A, S8B, and S8C). The unimodal distribution of sequence read counts per sample (sequencing depth) peaked at approximately 25,000 reads for *C. capistratus* and at 6,000 reads for *H. puella* (Figs. S9A and S9B). Non-metric multidimensional scaling (NMDS) of sequence read relative prey abundance data showed that *C. capistratus* from high coral cover reefs at the outer bay grouped together and were clearly separated from fish at the most degraded inner bay disturbed reefs (Fig. 4A). The diet composition of *H. puella* at the outer bay zone separated from both the inner bay and inner bay disturbed zones. However, there was no separation between the two zones located inside of the bay (Fig. 4B). Diet composition of *C. capistratus* was dominated by cnidarians at the outer bay high coral cover sites, whereas its diet was dominated by annelids at the inner bay disturbed zone with a more mixed diet at the inner bay zone (Fig. 4C, Fig. S10A). Within the phylum Cnidaria, fish at the outer bay preferentially fed upon hard corals in the family *Poritidae* together with soft corals (*Plexauridae*, *Gorgoniidae*, *Briareidae*), while anemones and *Porites* sp. were dominant diet items at the inner bay zone (Fig. S10B). In contrast, *Porites* sp. was less consumed at the inner bay disturbed zone, where sequence reads of both families *Mussidae* and *Merulinidae* and anemones (families: *Aiptasiidae*, *Boloceroididae, Discosomatidae*) were higher (Fig. S10B). Soft corals were present in negligible proportions in the diets of fish residing at both zones inside of the bay (Fig. 4C, Fig. S10A), while jellyfish appeared in the diet of fish in the inner bay disturbed zone (Fig. S11B). Across all zones there were low proportions of reads belonging to Corallimorpharia and Zoantharia (Fig. S10A).

**Figure 4.**
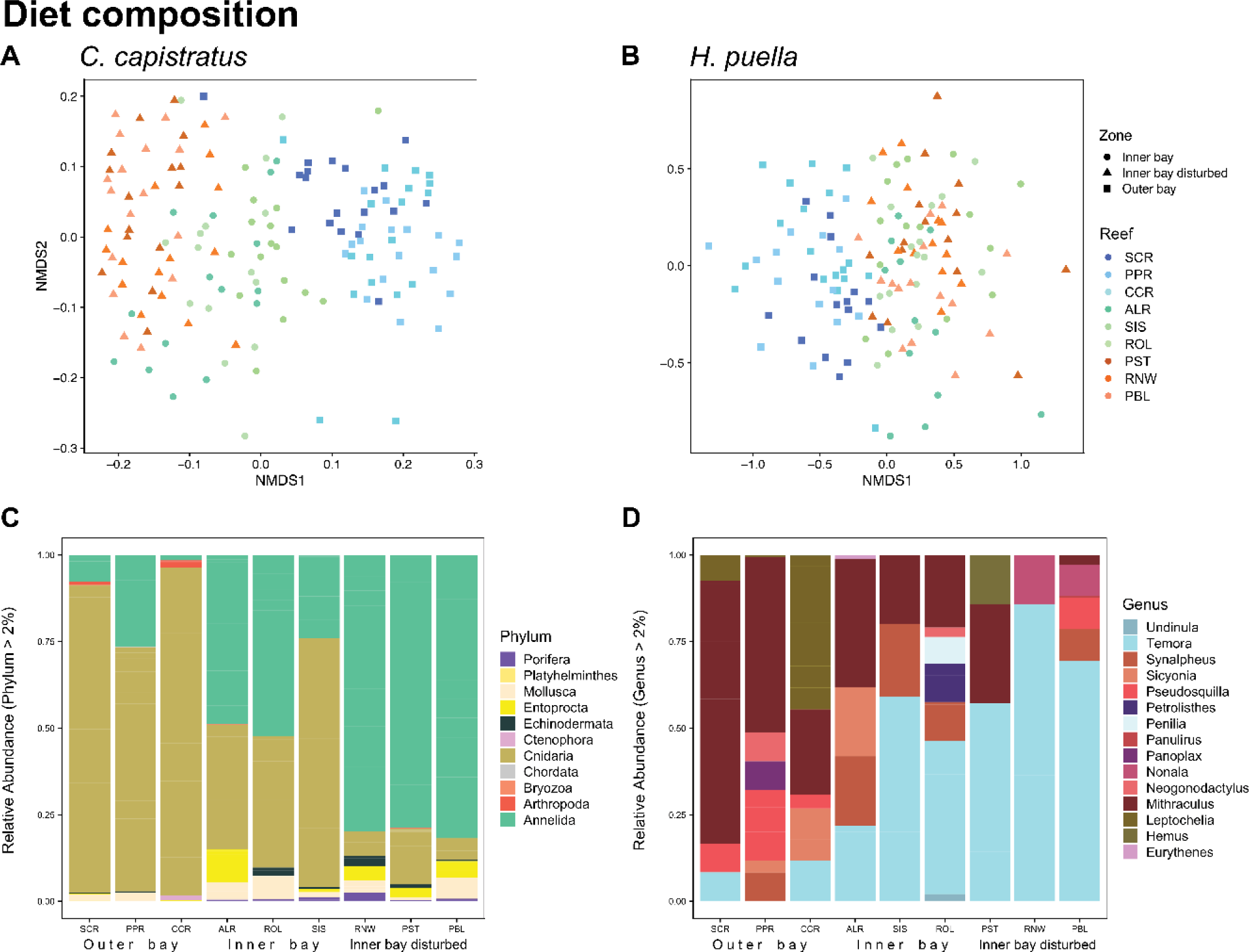
Differences in fish diet composition based on stomach and gut content metabarcoding across reefs. Nonmetric multidimensional scaling (NMDS) plots are based on Bray-Curtis distance matrices between individuals of (A) *Chaetodon capistratus* and (B) *Hypoplectrus puella*. Dots depict fish individuals; reef zones are coded by shapes and color: blue = outer bay, green = inner bay, and orange = inner bay disturbed. Variation in diet composition across the habitat gradient for (C) *C. capistratus* by phylum and (D) *H. puella* at the genus level within arthropods, their primary prey.

The diet of *H. puella* was dominated by arthropods at the phylum level across all reefs and zones (Fig. S11A). However, differences in diet composition among reefs and zones emerged at lower taxonomic levels. Within Arthropoda, more copepods were consumed inside of the bay and more decapods at the high coral cover outer bay reefs (Fig. S11B). When considering prey communities at the genus level, macrocrustaceans dominated the diet at outer bay reefs and microcrustaceans, many of them planktonic taxa, were prevalent in the diet across the inner bay and inner bay disturbed zones (Fig. 4D). At both zones located inside the bay, *H. puella*’s arthropod diet contained a large proportion of the copepod *Temora stylifera*, whereas diets at outer bay reefs were dominated by crabs in the genus *Mithraculus,* and to a lesser extent tanaid crustaceans (genus: *Leptochelia*) and mantis shrimp (genus: *Pseudosquilla*) (Fig. 4D), whereas prawns (genus: *Sicyonia*), rubble crabs (genus: *Panoplax*), snapping shrimp (genus: *Synalpheus*) and mantis shrimp (genus: *Neogonodactylus*) were consumed in smaller proportions (Fig. 4D). At the phylum level, Chordata constituted the second most important diet item of *H. puella* but relative read abundances were significantly lower than for Arthropoda (Fig.S11A). Chordates largely consisted of fishes (Fig. S12A), with Gobiformes being most frequently consumed followed by Blenniiformes (Fig. S12A). COI metabarcoding detected a broad taxonomic range of fishes (51 OTUs, class: Actinopterygii), of which 19 were identified at species level (Table 1). The chaenopsid blenny *Emblemariopsis arawak* was the most frequently detected species (at six of nine reefs) (Fig. S12B).

**Table 1:**
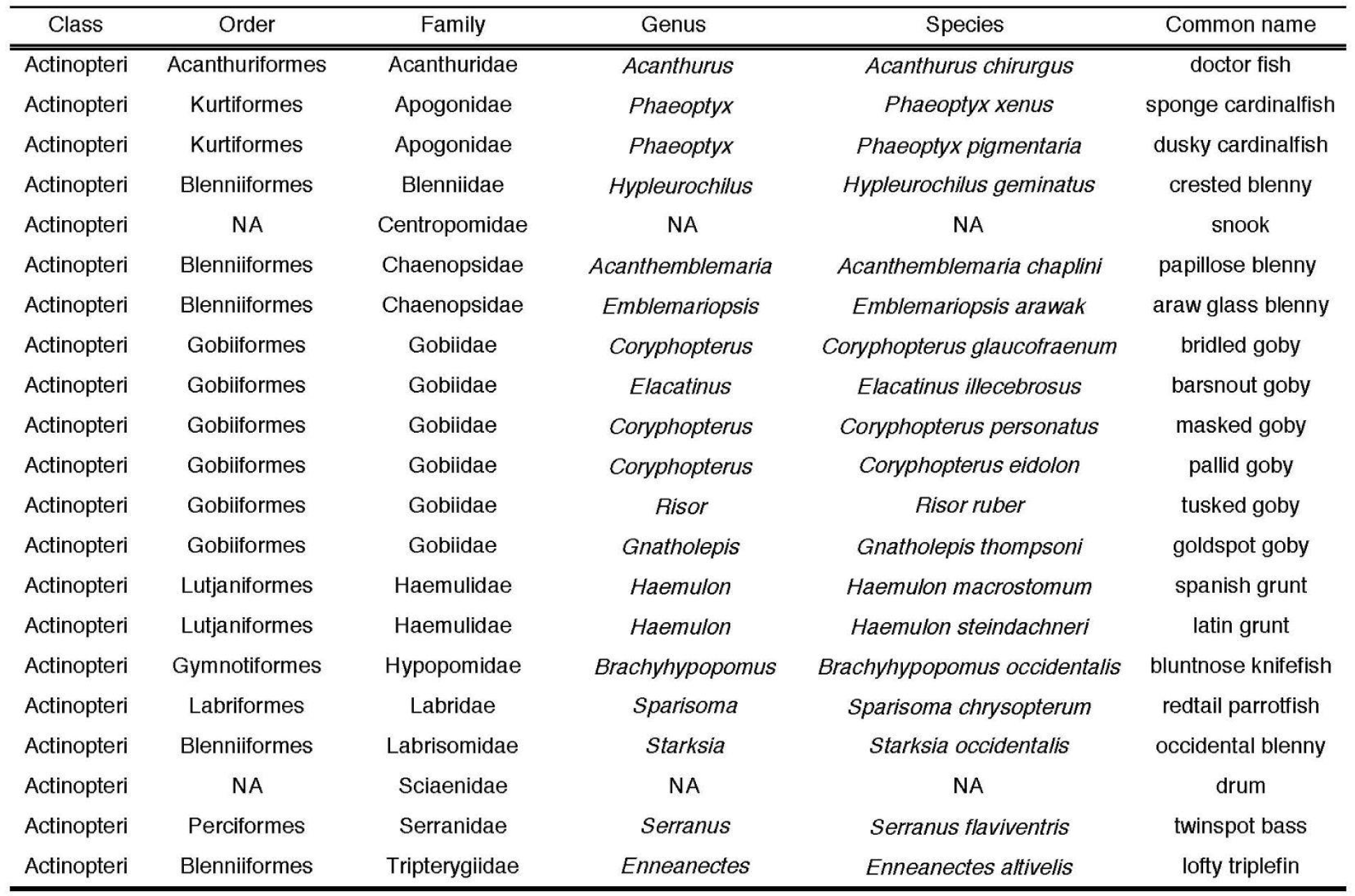
Fishes identified in the diet of *H. puella* (including only those OTUs that were identified to at least family level, 41% of all 51 OTUs assigned to Actinopteri).

### Relationship between coral cover and fish diet

Coral cover was a strong predictor of the relative abundance of hard corals and annelids in the diet of *C. capistratus* (Figs. 5A and 5B, Table 2). Reef zones varied in percent coral cover, and there was a significant relationship between the butterflyfish diet and zones (Table 2), with annelids decreasing and corals increasing in the diet when moving from inner bay disturbed zone to the inner bay zone and the outer bay zone. On the other hand, coral cover did not predict the relative abundance of the dominant prey of *H. puella*, benthic arthropods (Fig. 5C, Table 2), but it did predict consumption of planktonic arthropods (Fig. 5D, Table 2). We found no significant relationship between coral cover and the abundance of fishes detected within the guts of *H. puella* (Table 2). In addition, for both fish species we detected no significant effect of fish age on the proportions of main prey items in the diets (Table 2).

**Figure 5.**
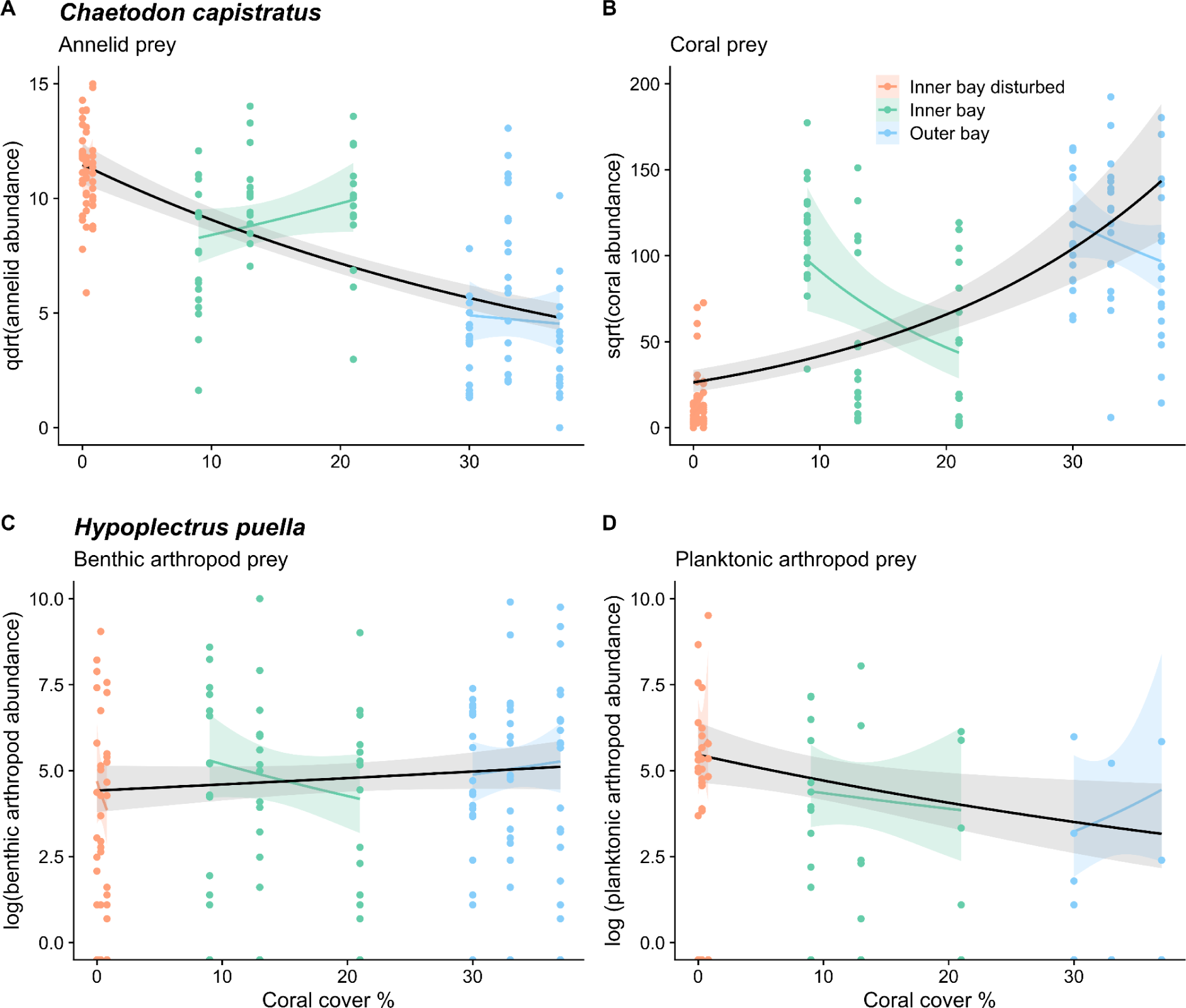
Coral cover effects on fish diets. Negative binomial generalized linear mixed-effects models (GLMMs) were fitted to predict the effect of percent coral cover on the sequencing read abundance of dominant diet items consumed by two fish species: *C. capistratus* feeding on (A) annelids and (B) hard coral, and *H. puella* feeding on (C) benthic arthropods and (D) planktonic arthropods. Smoothed black lines depict the overall trends across the coral cover gradient while coloured lines represent variability within each reef zone; the contrasting patterns between black and coloured lines suggest the presence of scale-dependent trends in prey consumption. Annelid read abundance was fourth root transformed, hard coral read abundance was square root transformed, and reads of both arthropod groups were log transformed to improve model fitting.

**Table 2.**
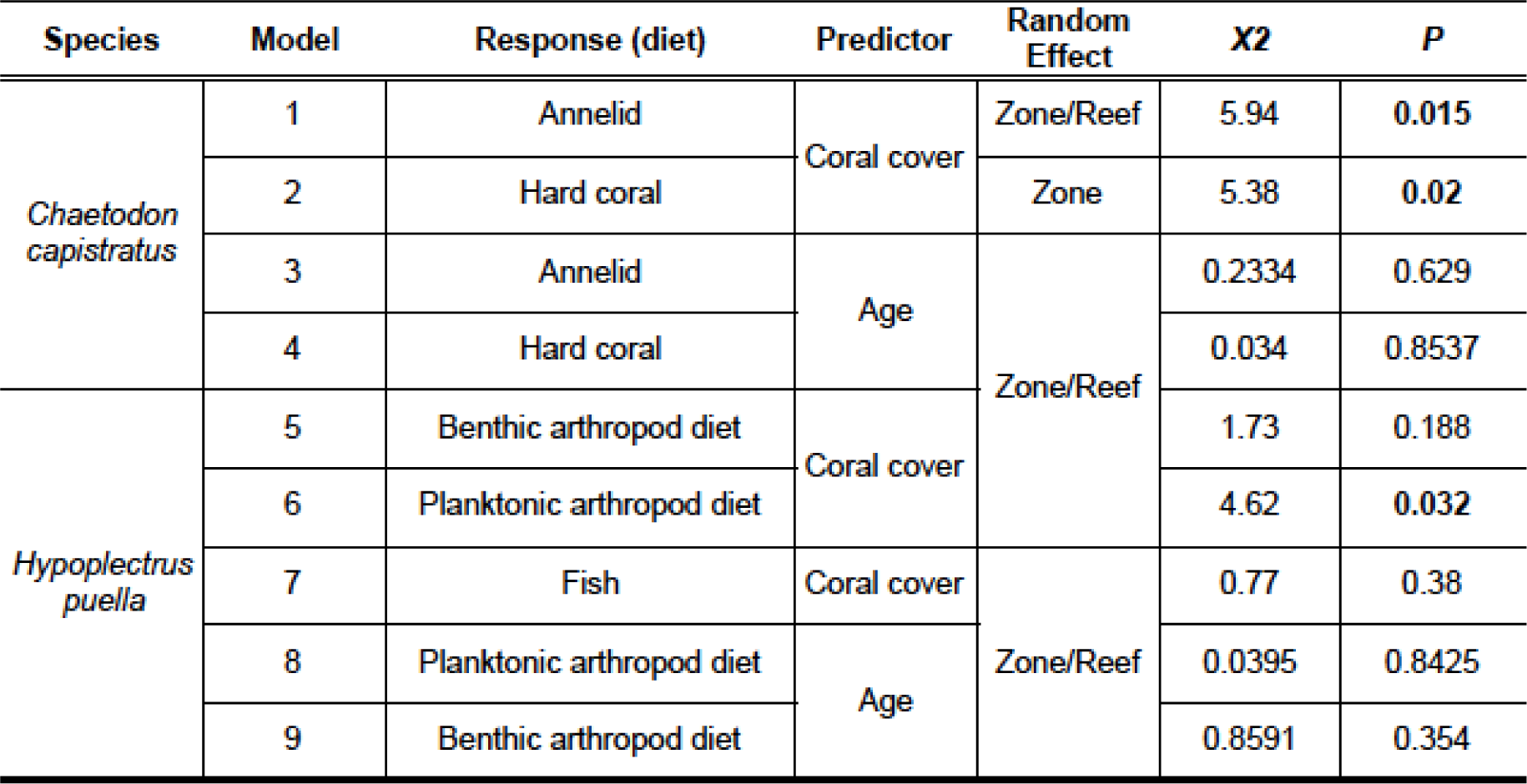
Results of general linear mixed effects models examining the effect of coral cover on different prey items in the diets of *Chaetodon capistratus* and *Hypoplectrus puella*. Coral cover accounted for variation in both the hard coral and annelid diet of *C. capistratus* and the consumption of pelagic arthropods in the diet of *H. puella.* Significant effects are depicted in bold.

### Diet strategy

Amundsen plots of fish diet strategy (Fig. 6A) across zones suggested that the diet of *C. capistratus* was dominated by very few prey items, as indicated by points located in the middle to upper right corner of the plots (Fig. 6B-D). This relatively specialized diet was complemented by a diverse array of occasional prey items that were consumed in low abundance (lower left corner of the plot). While *C. capistratus* appeared as a facultative specialist, *H. puella* displayed a generalist diet that was dominated by arthropods across all zones (Fig. 6E-G). Across the habitat gradient, *C. capistratus* switched its main diet item from hard coral, i.e., *Porites* sp. (phylum Cnidaria) at the outer bay zone (Fig. 6B-D), to a mix of *Porites* sp. and a sessile worm, *Loimia medusa* (phylum Annelida) at the inner bay zone (Fig. 6C), towards a diet dominated by *Loimia medusa* at the inner bay disturbed zone (Fig. 6D). The observed switch in the main diet item entailed that the diet of individual fish was less diverse (i.e., more specialized) as indicated by a decrease in the within phenotype component (WPC) at the disturbed zone in comparison with the other two zones (Fig. 6D). In the outer bay, *H. puella* consumed crabs in the genus *Mithraculus* frequently and in large quantities (25-50%) relative to other prey items, which were less apparent inside of the bay (<25%) (Fig. 6E-G). In contrast, the frequency of microcrustaceans in the diet was higher in inner bay zones, dominated by copepods in the genus *Temora* (Fig. 6F and G). At both inner bay zones, we found that the diet among individuals was more variable as indicated by an increase in the between phenotype component (BPC), implying an increase in individual specialization and a broader diet on the population level at these reefs (Fig. 6F and G) relative to outer bay reefs (Fig. 6E).

**Figure 6.**
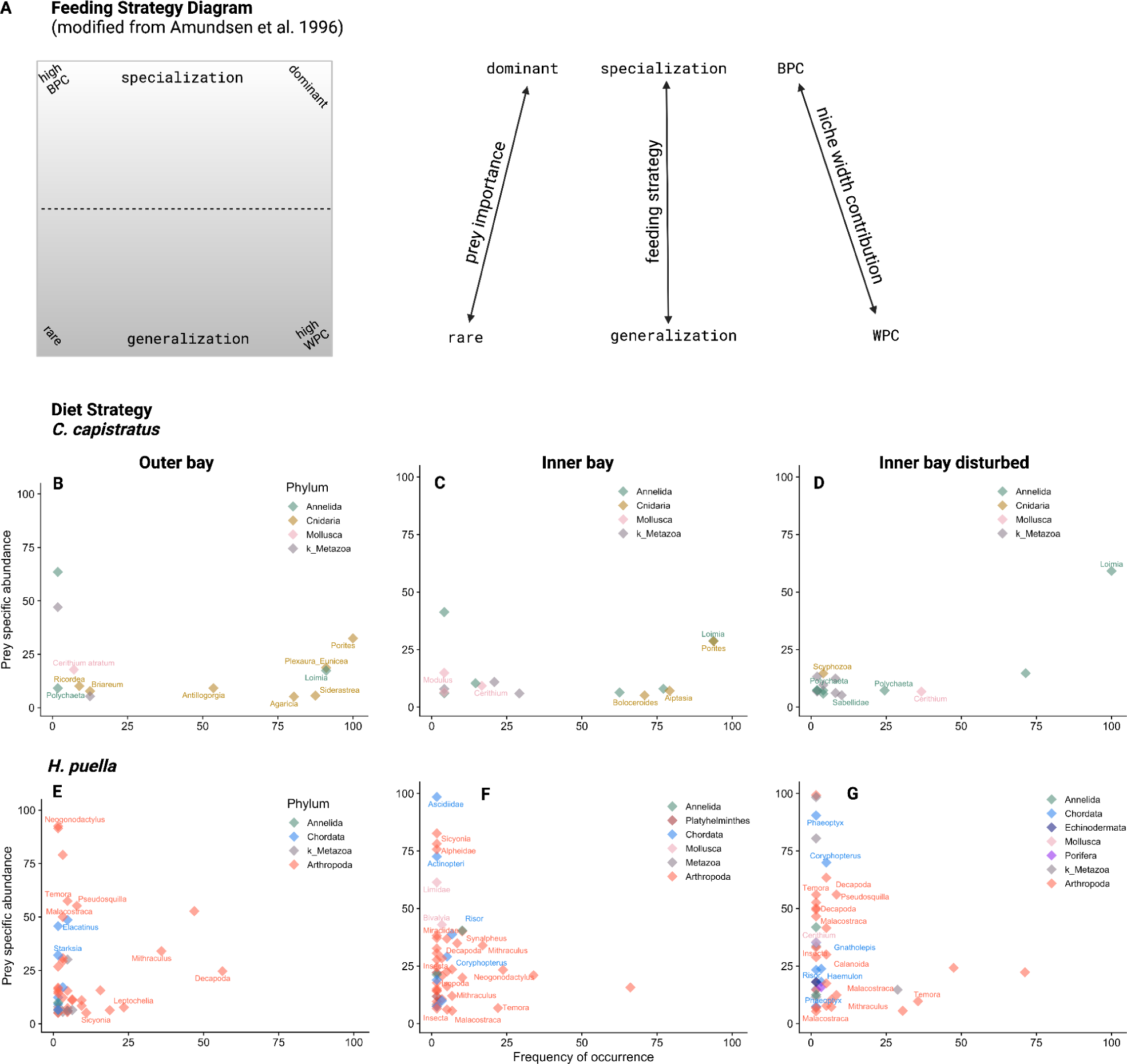
Fish feeding strategies across the habitat gradient. (A) Schematic diagram modified from Amundsen et al. (1996) illustrating how feeding strategy as shaped by niche width contribution and prey importance is inferred from i) the vertical axes indicating specialization (upper portion of plot) and generalization (lower portion of plot) and ii) the diagonal axis representing the Within Phenotype Component (WPC, lower right corner) and Between Phenotype Component (BPC, upper left corner) indices. (B-G) Graphical analysis of fish diet strategy at three reef zones using relative read abundance data for *C. capistratus* (B,C,D), and *H. puella* (E,F,G) following the method of Amundsen et al. (1996) based on and modified from Costello (1990). Points represent OTUs assigned to prey taxa. Points located in the upper right indicate a specialized diet at the population level (abundant and frequent diet items), whereas points in the lower left corner indicate opportunistic, occasional diet items that are found rarely and in low abundance in the diet. Frequency of occurrence = percentage of fish in which a prey item was present versus the total number of fish. Prey specific abundance = percentage of diet made-up by a given prey item (OTU) across only the number of fish individuals where it occurred.

## Discussion

Our study was conducted across three habitat zones featuring different levels of coral diversity and cover that represented degrees of reef degradation associated with variation in associated invertebrate assemblages. Against this backdrop we showed that the diet composition of two benthic-feeding coral reef fish species (*C. capistratus* and *H. puella*) with distinct feeding strategies was influenced by the proportion of live coral cover. Combining ecological data, high-resolution diet data and otolith analysis revealed relationships between benthic cover, resource use, and fish growth and body condition. We found dietary differences in both fish species between healthy and degraded reefs; however, the extent to which dietary adjustments shield potentially adverse effects of spatial differences in prey availability varied between fish species, as evidenced by reduced growth and body condition observed only in the browsing species inhabiting the most degraded reefs. Our results suggest that fish trophic roles can vary within species across small spatial scales in relation.

### Changes in fish diet with reef degradation

Contrasting our expectation, *C. capistratus* did not broaden its diet where coral cover was low, but switched preference while narrowing its diet. It maintained its browsing feeding mode (this is, moving between resource patches nipping on sessile prey) on degraded reefs by primarily feeding on terebellid worms (phylum: Annelida, class: Polychaeta) resembling the coral polyps that they typically consume in providing an evenly distributed and abundant prey resource. This feeding behavior potentially allows individuals to maintain a similar energy budget between healthy and degraded reefs (Uchida et al., 2007; Van Leeuwen et al., 2013). While *C. capistratus* has been previously reported to include polychaete tentacles in its diet (e.g., families: Serpulidae, Terebellidae), the percent volume detected in stomachs was commonly low in relation to cnidarian prey of largely hexacorals and octocorals. For example, Birkeland and Neudecker (1981) showed that *C. capistratus* can complement its anthozoan dominated diet (80%) with items of high nutritional value such as polychaete worms (Rotjan & Lewis, 2008), but no previous study has shown a near complete switch. Prey may need to exceed a certain abundance threshold to represent a diet item worth exploiting for *C. capistratus,* and previous studies from the 1980s were conducted on much less disturbed reefs. Our findings are consistent with previous studies finding *C. capistratus* to feed selectively (Birkeland & Neudecker, 1981; Gore, 1984; Lasker, 1985; Liedke et al. 2018; Casement 2021). *Chaetodon capistratus* specialized on chemically defended worm tentacles at degraded reefs, indicating an adaptation to feeding on chemically defended prey. This dietary specialization may serve to avoid competition with the Caribbean congeners *C. striatus* and *C. ocellatus*, which are known to be less tolerant of dietary allelochemicals (Pitts, 1991; Vrolijk et al., 1995; Liedke et al., 2018), thus highlighting a potential ecological advantage in resource utilization within degraded reef environments.

*Hypolplectrus puella* likely fed in relation to resource availability as both the consumption and relative density of decapod crustaceans decreased at the inner bay disturbed zone. In addition, *H. puella’*s dietary preference differed across the habitat gradient as our data suggest that it actively avoided a particular diet item (mithracid crabs; infraorder: Brachyura) on degraded reefs while increasingly consuming copepods. Despite their small size, copepods potentially provide a more numerous and evenly distributed food source than crabs under degraded conditions. However, as calanoid copepod distribution has been shown to be uniform across our study area (Rodas et al., 2020), the observed dietary pattern was likely not driven by relative plankton availability, but rather changes in the accessibility of macrocrustacean prey. In contrast to the different crustacean prey (macro vs microcrustaceans), the contribution of fishes in the diet of *H. puella*, which constituted the second most common diet item, did not vary with coral cover. Overall, the relative proportion of fish prey consumed by *H. puella* resembled that of its congener *H. unicolor* (the butter hamlet) at our study area, and was thus higher than previously reported (Puebla et al., 2018). Our metabarcoding approach likely surpassed previous visual analyses that might have underestimated the proportion and diversity of fish prey in the diet of *H. puella* (∼10%, Randall, 1967; Whiteman, Côté, and Reynolds 2007; Puebla et al., 2018) as fish can be digested faster than crustaceans (e.g., four times faster in rock cod, Beukers-Stewart & Jones, 2004). In addition, *Hypoplectrus puella* targeted fish species that were previously not detected in *H. unicolor’s* diet at Bocas del Toro (except for *Coryphopterus personatus*) (Puebla et al., 2018), suggesting that these two species might partition their fish prey to some extent. *Hypoplectrus puella* also consumed proportionally more crabs than *H. unicolor*, further indicating dietary partitioning. These findings contrast with previous diet analyses reporting high levels of dietary overlap among hamlet species suggestive of ecological equivalence (Whiteman, Côté, and Reynolds 2007; Holt et al., 2008).

### Changes in fish condition with reef degradation

Our results suggest that low coral cover reefs potentially provide less suitable resources for *C. capistratus*, whereas *H. puella* appears more resilient. We found optimal body condition levels (median Kn ≥ 1) in *C. capistratus* at both the inner bay and outer bay zones, suggesting that both high and intermediate levels of live coral cover provide favorable foraging grounds for this species, as opposed to degraded reefs. In contrast, body condition was significantly lower in medium sized individuals at the inner bay disturbed zone, where they also grew significantly slower. Together these results suggest that the annelid diet at low coral cover reefs was suboptimal in comparison to a mixed or coral-dominated diet. Mixed diets have been proposed to enhance fitness (balanced diet hypothesis, Pulliam, 1975); however; recent meta-analysis found the ‘single optimal prey item’ better promoted predator fitness (Lefcheck et al., 2013). Yet only the largest fish seemed to fare well on a diet that was dominated by one prey item on degraded reefs. Additionally, *C. capistratus* individuals from the inner bay zone were significantly older than those caught on reefs in the other zones. This suggests that survival rates were higher in this zone, likely due to the combination of sufficient live coral cover levels in comparison to the inner bay disturbed zone, and potentially lower predation pressure than at the outer bay zone where coral cover was highest and fishing pressure likely lower potentially implying higher predation pressure on butterflyfishes (Cramer, 2013; Seemann et al., 2018). Conspicuously, the density of *C. capistratus* did not significantly vary among zones and was thus decoupled from variation in body condition. The population size of versatile feeders may only slowly respond to changes in prey availability; for example, the most specialized species of Indo-Pacific butterflyfishes, but not the more generalist species, were shown to have population sizes limited by resource availability (Lawton & Pratchett, 2012). Some of the few previous studies examining the effects of coral reef habitat on fish body condition found reduced levels associated with dietary changes (Pratchett et al., 2004; Berumen et al., 2005; Hempson et al., 2017). Although *C. capistratus* densities were stable despite reduced body condition at degraded reefs, our results highlight that long-term sublethal effects of habitat degradation may manifest over time.

Our data suggest that the condition of *H. puella* did not decrease with decreasing coral cover despite the marked difference in size between its crustacean prey across the habitat gradient, demonstrating a versatile nutritional and behavioral physiology. The size ratio between fish and its prey influences the effort needed for searching prey and the relative contribution of a prey to a predator’s energy needs (Hart & Gill, 1993). Feeding on planktonic prey suggests low search effort but also low caloric return per prey (Hart & Gill, 1993). However, the increased reliance of *H. puella* on an alternative food source providing lower nutritional value (Hart & Gill, 1993) had no negative effect on its mean density, body condition or growth rate, suggesting the ease of obtaining plankton made-up for its relatively low source of energy. Overall, in contrast to previous results showing that shifted diets on degraded reefs led to less energy stored in the livers of another coral reef mesopredator (Hempson et al., 2018), our results suggest that *H. puella* successfully coped with degraded reef conditions.

### Implications for management and conservation

While most pronounced in *C. capistratus*, both species differed in their trophic functions across the habitat gradient facilitated by flexible feeding behavior, which may shape food webs in different ways at healthy versus degraded reefs. The predominant prey items of *C. capistratus* and *H. puella* at high coral cover reefs (live corals and macrocrustaceans, respectively) rely upon planktonic carbon sources and symbiotic photosynthesis in the case of corals, and epibenthic food in the case of many crustaceans. At degraded reefs in contrast, terebellid worms and calanoid copepods use detrital deposits and phytoplankton, respectively. This implies that both fish species used different trophic pathways at healthy and degraded reefs. Habitat degradation in the form of fragmentation and land use change has previously led to a simplified food web structure in tidal creeks in the Bahamas and a freshwater system in Croatia (Layman et al., 2007; Price et al., 2019). The reliance of *H. puella* on planktonic food in the disturbed zone was in line with previous findings suggesting that pelagic food sources may increasingly support coral reef fishes on degraded reefs (Morais & Bellwood, 2019). Further investigation of prey diets or verification with other trophic markers (e.g., compound-specific stable isotopes; McMahon et al., 2016) could further substantiate carbon sources.

Our study adds to recent work on how consumer-resource interactions alter coral reef food webs in response to degradation at relatively small spatial scales (Layman et al., 2007; Karkarey et al., 2017; Hempson et al., 2018; Semmler et al., 2022), thereby influencing energy flow and ultimately ecosystem functioning (Duffy et al., 2007). Species interactions are thought to underpin stabilizing mechanisms, such as functional redundancy (Rosenfeld, 2002) and trophic compensation (e.g., Ghedini, Russell, and Connell, 2015), under conditions of environmental change. Yet this notion is being increasingly scrutinized in the case of coral reefs, where high levels of niche partitioning may render coral reef fish assemblages more vulnerable than previously assumed (Brandl & Bellwood, 2014; Kramer et al., 2015; Leray, Meyer and Mills, 2015; Bejarano et al., 2017; Brandl, Casey and Meyer, 2020; Semmler et al., 2021). Intraspecific dietary variation as documented by the present study may play a role in mediating this pattern (Albert et al., 2010) and thus potentially affect levels of functional redundancy within the fish assemblage at a given location. In addition, our findings question the usefulness of coarse trophic classifications of species that largely lack empirical ground-truthing. For example, Parravicini et al. (2020) found coral reef fish invertivores were more diversified than suggested by previous classifications. Our study corroborates their findings and improves the resolution of benthic invertivore diets on coral reefs.

Our finding that *C. capistratus* exhibited reduced growth at degraded reefs and greater dietary variation among reef zones in contrast to *H. puella*, suggests that benthic invertivorous fishes that specialize on sessile taxa may be more susceptible to reef degradation than those that specialize on free-living invertebrates. High resolution DNA-based analysis revealed that within-species dietary variation was greater than previously thought for our study species. We further demonstrated that versatile feeding behavior can entail the use of different trophic pathways between high and low coral cover reefs. Consequently, we advocate taking into account diet versatility and limitations on body condition in putatively generalist strategies, as they can influence both species persistence and tropho-dynamics on degrading coral reefs.

## Acknowledgements

We thank Lucía Rodriguez for field assistance, Joan Antaneda for her help in the laboratory, Ross Whippo for conducting the fish survey, and Clare Fieseler for taking photos of the benthos. The staff of the Bocas del Toro Research Station provided logistical support. We are grateful to Kristin Saltonstall and Marta Vargas for their support at the Smithsonian Tropical Research Institute’s (STRI) Ecological and Evolutionary Genomics Laboratory. We thank Justin Touchon for guidance with mixed models, Rainer Froese for advice on fish length-weight relationships and Marco Smolla for support with R functions. Friederike Clever was supported by a Smithsonian Short Term Fellowship and a PhD studentship by Manchester Metropolitan University. A research permit was issued by the Ministerio de Ambiente Panamá (No. SE/A-113-17).

## Author contributions statement

F.C., M.L. and R.F.P. conceived and designed the study. F.C. and M.L. conducted the fieldwork. F.C. dissected the fish stomachs and guts and extracted the DNA. F.C. and M.L. prepared the DNA for sequencing. B.N. processed the sequencing data. B.D.G., A.O’D. and H.Q. conducted the otolith analysis. R.F.P., N.K., A.H.A., W.O.M., A.O’D. and M.L. contributed reagents and supplies. F.C. analyzed the data and wrote the first draft of the manuscript with input from M.L., R.F.P., and A.H.A. B.D.G. and A.O’D. contributed to a later version of the manuscript. All authors reviewed the manuscript and contributed to the final version.

## Data and code availability

Sequencing data has been submitted to the NCBI Short Read Archive (SRA) database (https://www.ncbi.nlm.nih.gov/sra) under bioproject number Accession:XXXXX. Raw data and R code are available on Dryad Digital Repository XXXX.

## References

Alberdi, A., Aizpurua, O., Bohmann, K., Gopalakrishnan, S., Lynggaard, C., Nielsen, M., & Gilbert, M. T. P. (2019). Promises and pitfalls of using high-throughput sequencing for diet analysis. Molecular Ecology Resources, 19(2), 327–348. 10.1111/1755-0998.12960

Alberdi, A., Aizpurua, O., Gilbert, M. T. P., & Bohmann, K. (2017). Scrutinizing key steps for reliable metabarcoding of environmental samples. Methods in Ecology and Evolution. 10.1111/2041-210X.12849

Albert, C. H., Thuiller, W., Yoccoz, N. G., Douzet, R., Aubert, S., & Lavorel, S. (2010). A multi-trait approach reveals the structure and the relative importance of intra-vs. Interspecific variability in plant traits. Functional Ecology, 24(6), 1192–1201. 10.1111/j.1365-2435.2010.01727.x

Altieri, A. H., Harrison, S. B., Seemann, J., Collin, R., Diaz, R. J., & Knowlton, N. (2017). Tropical dead zones and mass mortalities on coral reefs. Proceedings of the National Academy of Sciences of the United States of America, 114(14), 3660–3665. 10.1073/pnas.1621517114

Amundsen, P. A., Gabler, H. M., & Staldvik, F. J. (1996). A new approach to graphical analysis of feeding strategy from stomach contents data-modification of the Costello (1990) method. Journal of Fish Biology, 48(4), 607–614. 10.1111/j.1095-8649.1996.tb01455.x

Anderson, W., Carpenter, K. E., Gilmore, G., Milagrosa Bustamante, G., & Robertson, R. (2015). Hypoplectrus puella (Barred Hamlet). The IUCN Red List of Threatened Species 2015. https://www.iucnredlist.org/species/16759126/16781798

Baker, R., Buckland, A., & Sheaves, M. (2014). Fish gut content analysis: Robust measures of diet composition. Fish and Fisheries, 15(1), 170–177. 10.1111/faf.12026

Barlow, G. W. (1975). On the sociobiology of some hermaphroditic serranid fishes, the hamlets, in Puerto Rico. Marine Biology, 33(4), 295–300. 10.1007/BF00390567

Bates, D., Mächler, M., Bolker, B. M., & Walker, S. C. (2015). Fitting linear mixed-effects models using lme4. Journal of Statistical Software, 67(1). 10.18637/jss.v067.i01

Bejarano, S., Jouffray, J. B., Chollett, I., Allen, R., Roff, G., Marshell, A., Steneck, R., Ferse, S. C. A., & Mumby, P. J. (2017). The shape of success in a turbulent world: Wave exposure filtering of coral reef herbivory. Functional Ecology, 31(6), 1312–1324. 10.1111/1365-2435.12828

Benjamini, Y., & Hochberg, Y. (1995). Controlling the false discovery rate: A practical and powerful approach to multiple testing. Journal of the Royal Statistical Society: Series B (Methodological), 57(1), 289–300. 10.1111/j.2517-6161.1995.tb02031.x

Berry, O., Bulman, C., Bunce, M., Coghlan, M., Murray, D. C., & Ward, R. D. (2015). Comparison of morphological and DNA metabarcoding analyses of diets in exploited marine fishes. Marine Ecology Progress Series, 540, 167–181. 10.3354/meps11524

Berumen, M. L., Pratchett, M. S., & McCormick, M. I. (2005). Within-reef differences in diet and body condition of coral-feeding butterflyfishes (Chaetodontidae). Marine Ecology Progress Series, 287, 217–227. 10.3354/meps287217

Beukers-Stewart, B. D., & Jones, G. P. (2004). The influence of prey abundance on the feeding ecology of two piscivorous species of coral reef fish. Journal of Experimental Marine Biology and Ecology, 299(2), 155–184. 10.1016/j.jembe.2003.08.015

Birkeland, C. & Neudecker, S. (1981). Foraging behavior of two Caribbean chaetodontids: *Chaetodon capistratus* and *C. aculeatus*. Copeia, 1981(169–178).

Blackwell, B. G., Brown, M. L., & Willis, D. W. (2000). Relative weight (Wr) status and current Use in fisheries assessment and management. Reviews in Fisheries Science, 8(1), 1–44. 10.1080/10641260091129161

Bolnick, D. I., Svanbäck, R., Fordyce, J. A., Yang, L. H., Davis, J. M., Hulsey, C. D., & Forister, M. L. (2003). The ecology of individuals: Incidence and implications of individual specialization. American Naturalist, 161(1), 1–28. 10.1086/343878

Brandl, S. J., & Bellwood, D. R. (2014). Individual-based analyses reveal limited functional overlap in a coral reef fish community. Journal of Animal Ecology, 83(3), 661–670. 10.1111/1365-2656.12171

Brandl, S. J., Casey, J. M., & Meyer, C. P. (2020). Dietary and habitat niche partitioning in congeneric cryptobenthic reef fish species. Coral Reefs. 10.1007/s00338-020-01892-z

Bray, J. R., & Curtis, J. T. (1957). An ordination of the upland forest communities of southern Wisconsin. Ecological Monographs, 27(4), 325–349. 10.2307/1942268

Callahan, B. J., McMurdie, P. J., Rosen, M. J., Han, A. W., Johnson, A. J. A., & Holmes, S. P. (2016). DADA2: High-resolution sample inference from Illumina amplicon data. Nature Methods, 13, 581–581.

Casement, E. A. (2021). Abundance, foraging levels, and dietary preferences of Chaetodon capistratus on reefs surrounding Porvenir Island in the Guna Yala Comarca of Panamá. Independent Study Project (ISP) Collection. 3620. https://digitalcollections.sit.edu/isp_collection/3620

Chariton, A. A., Stephenson, S., Morgan, M. J., Steven, A. D. L., Colloff, M. J., Court, L. N., & Hardy, C. M. (2015). Metabarcoding of benthic eukaryote communities predicts the ecological condition of estuaries. Environmental Pollution, 203, 165–174.

Clarke, K. R., & Warwick, R. M. (2001). Change in Marine Communities: An Approach to Statistical analysis and Interpretation, 2, 1–168

Clever, F., Sourisse, J. M., Preziosi, R. F., Eisen, J. A., Guerra, E. C. R., Scott, J. J., Wilkins, L. G. E., Altieri, A. H., McMillan, W. O., & Leray, M. (2022). The gut microbiome variability of a butterflyfish increases on severely degraded Caribbean reefs. Communications Biology, 5(1), 1–15. 10.1038/s42003-022-03679-0

Coker, D. J., DiBattista, J. D., Stat, M., Arrigoni, R., Reimer, J., Terraneo, T. I., Villalobos, R., Nowicki, J. P., Bunce, M., & Berumen, M. L. (2022). DNA metabarcoding confirms primary targets and breadth of diet for coral reef butterflyfishes. Coral Reefs. 10.1007/s00338-022-02302-2

Collin, R., D’Croz, L., Gondola, P., & Del Rosario, J. B. 2009. Climate and hydrological factors affecting variation in chlorophyll concentration and water clarity in the Bahia Almirante, Panama. in Proceedings of the Smithsonian Marine Science Symposium, edited by Lang, Michael A., MacIntyre, Ian G., & Ruetzler, Klaus., First ed. 323–334. Smithsonian Contributions to the Marine Sciences. Washington DC: Smithsonian Institution Scholarly Press. 10.5479/10088/19175

Costello, M. J. (1990). Predator feeding strategy and prey importance: A new graphical analysis. Journal of Fish Biology, 36(2), 261–263. 10.1111/j.1095-8649.1990.tb05601.x

Cramer, K. L. (2013). History of Human Occupation and Environmental Change in Western and Central Caribbean Panama. Bulletin of Marine Science, 89(4), 955–982. 10.5343/bms.2012.1028

D’Croz, L., Rosario, J. B. del, & Gondola, P. (2005). The effect of fresh water runoff on the distribution of dissolved inorganic nutrients and plankton in the Bocas Del Toro Archipelago, Caribbean Panamá. Caribbean Journal of Science, 41(3), 414–429.

De Barba, M., Miquel, C., Boyer, F., Mercier, C., Rioux, D., Coissac, E., & Taberlet, P. (2014). DNA metabarcoding multiplexing and validation of data accuracy for diet assessment: Application to omnivorous diet. Molecular Ecology Resources, 14(2), 306–323. 10.1111/1755-0998.12188

Donelson, J. M., Sunday, J. M., Figueira, W. F., Gaitán-Espitia, J. D., Hobday, A. J., Johnson, C. R., Leis, J. M., Ling, S. D., Marshall, D., Pandolfi, J. M., Pecl, G., Rodgers, G. G., Booth, D. J., & Munday, P. L. (2019). Understanding interactions between plasticity, adaptation and range shifts in response to marine environmental change. Philosophical Transactions of the Royal Society B: Biological Sciences, 374(1768). 10.1098/rstb.2018.0186

Doucette, V. E., Rodriguez Bravo, L. M., Altieri, A. H., & Johnson, M. D. (2022). Negative effects of a zoanthid competitor limit coral calcification more than ocean acidification. Royal Society Open Science, 9(11), 220760. 10.1098/rsos.220760

Duffy, J. E., Cardinale, B. J., France, K. E., McIntyre, P. B., Thébault, E., & Loreau, M. (2007). The functional role of biodiversity in ecosystems: Incorporating trophic complexity. Ecology Letters, 10(6), 522–538. 10.1111/j.1461-0248.2007.01037.x

Dunn, O. J. (1964). Multiple comparisons using rank sums. Technometrics, 6(3), 241–252. 10.1080/00401706.1964.10490181

Froese, R. (2006). Cube law, condition factor and weight-length relationships: History, meta-analysis and recommendations. Journal of Applied Ichthyology, 22(4), 241–253. 10.1111/j.1439-0426.2006.00805.x

Frøslev, T. G., Kjøller, R., Bruun, H. H., Ejrnæs, R., Brunbjerg, A. K., Pietroni, C., & Hansen, A. J. (2017). Algorithm for post-clustering curation of DNA amplicon data yields reliable biodiversity estimates. Nature Communications, 8(1), 1–11. 10.1038/s41467-017-01312-x

Gao, X., Lin, H., Revanna, K., & Dong, Q. (2017). A Bayesian taxonomic classification method for 16S rRNA gene sequences with improved species-level accuracy. BMC Bioinformatics, 18(1). 10.1186/s12859-017-1670-4

Ghedini, G., Russell, B. D., & Connell, S. D. (2015). Trophic compensation reinforces resistance: herbivory absorbs the increasing effects of multiple disturbances. Ecology Letters, 18(2), 182–187.

Geller, J., Meyer, C., Parker, M., & Hawk, H. (2013). Redesign of PCR primers for mitochondrial cytochrome c oxidase subunit I for marine invertebrates and application in all-taxa biotic surveys. Molecular Ecology Resources, 13(5), 851–861. 10.1111/1755-0998.12138

Gochfeld, D. J. (2004). Predation-induced morphological and behavioral defenses in a hard coral: Implications for foraging behavior of coral-feeding butterflyfishes. Marine Ecology Progress Series, 267, 145–158. 10.3354/meps267145

Gore, M. A. (1984). Factors affecting the feeding behavior of a coral reef fish, *Chaetodon capistratus*. Bulletin of Marine Science, 35(2), 211–220.

Graham, N. A. J. (2007). Ecological versatility and the decline of coral feeding fishes following climate driven coral mortality. Marine Biology, 153(2), 119–127. 10.1007/s00227-007-0786-x

Hart, P. J. B., & Gill, A. B. (1993). Choosing prey size: A comparison of static and dynamic foraging models for predicting prey choice by fish. Marine Behaviour and Physiology, 23(1–4), 91–104. 10.1080/10236249309378859

Hempson, T. N., Graham, N. A. J., MacNeil, M. A., Bodin, N., & Wilson, S. K. (2018). Regime shifts shorten food chains for mesopredators with potential sublethal effects. Functional Ecology, 32(3), 820–830. 10.1111/1365-2435.13012

Hempson, T. N., Graham, N. A. J., MacNeil, M. A., Williamson, D. H., Jones, G. P., & Almany, G. R. (2017). Coral reef mesopredators switch prey, shortening food chains, in response to habitat degradation. Ecology and Evolution, 7(8), 2626–2635. 10.1002/ece3.2805

Holt, B. G., Emerson, B. C., Newton, J., Gage, M. J. G., & Côté, I. M. (2008). Stable isotope analysis of the *Hypoplectrus* species complex reveals no evidence for dietary niche divergence. Marine Ecology Progress Series, 357, 283–289. 10.3354/meps07339

Hyslop, E. J. (1980). Stomach contents analysis—A review of methods and their application. Journal of Fish Biology, 17(4), 411–429. 10.1111/j.1095-8649.1980.tb02775.x

Karkarey, R., Alcoverro, T., Kumar, S., & Arthur, R. (2017). Coping with catastrophe: Foraging plasticity enables a benthic predator to survive in rapidly degrading coral reefs. Animal Behaviour, 131, 13–22. 10.1016/j.anbehav.2017.07.010

Kaufmann, K. W., & Thompson, R. C. (2005). Water temperature variation and the meteorological and hydrographic environment of Bocas del Toro, Panama. Caribbean Journal of Science, 41, 392–413.

Kramer, M. J., Bellwood, O., Fulton, C. J., & Bellwood, D. R. (2015). Refining the invertivore: Diversity and specialization in fish predation on coral reef crustaceans. Marine Biology, 162(9), 1779–1786. 10.1007/s00227-015-2710-0

Kruskal, W. H., & Wallis, W. A. (1952). Use of ranks in one-criterion variance analysis. Journal of the American Statistical Association, 47(260), 583–621. 10.1080/01621459.1952.10483441

Lasker, H. R. (1985). Prey preferences and browsing pressure of the butterflyfish *Chaetodon capistratus* on Caribbean gorgonians. Marine Ecology Progress Series, 21, 213–220. 10.3354/meps021213

Lawton, R. J., & Pratchett, M. S. (2012). Influence of dietary specialization and resource availability on geographical variation in abundance of butterflyfish. Ecology and Evolution, 2(7), 1347–1361. 10.1002/ece3.253

Layman, C. A., Quattrochi, J. P., Peyer, C. M., & Allgeier, J. E. (2007). Niche width collapse in a resilient top predator following ecosystem fragmentation. Ecology Letters, 10(10), 937–944. 10.1111/j.1461-0248.2007.01087.x

Le Cren, E. D. (1951). The length-weight relationship and seasonal cycle in gonad weight and condition in the perch (*Perca fluviatilis*). The Journal of Animal Ecology, 20(2), 201–201. 10.2307/1540

Lefcheck, J. S., Whalen, M. A., Davenport, T. M., Stone, J. P., & Duffy, J. E. (2013). Physiological effects of diet mixing on consumer fitness: A meta-analysis. Ecology, 94(3), 565–572. 10.1890/12-0192.1

Leray, M., Agudelo, N., Mills, S. C., & Meyer, C. P. (2013b). Effectiveness of annealing blocking primers versus restriction enzymes for characterization of generalist diets: Unexpected prey revealed in the gut contents of two coral reef fish species. PLoS One, 8(4), e58076–e58076. 10.1371/journal.pone.0058076

Leray, M., Alldredge, A. L., Yang, J. Y., Meyer, C. P., Holbrook, S. J., Schmitt, R. J., Knowlton, N., & Brooks, A. J. (2019). Dietary partitioning promotes the coexistence of planktivorous species on coral reefs. Molecular Ecology, 28(10), 2694–2710. 10.1111/mec.15090

Leray, M., & Knowlton, N. (2015). DNA barcoding and metabarcoding of standardized samples reveal patterns of marine benthic diversity. Proceedings of the National Academy of Sciences U S A, 112(7), 2076–2081. 10.1073/pnas.1424997112

Leray, M., Knowlton, N., & Machida, R. J. (2022). MIDORI2: A collection of quality controlled, preformatted, and regularly updated reference databases for taxonomic assignment of eukaryotic mitochondrial sequences. Environmental DNA, 4(4), 894–907. 10.1002/edn3.303

Leray, M., Meyer, C. P., & Mills, S. C. (2015). Metabarcoding dietary analysis of coral dwelling predatory fish demonstrates the minor contribution of coral mutualists to their highly partitioned, generalist diet. PeerJ, 3, e1047–e1047. 10.7717/peerj.1047

Leray, M., Wilkins, L. G. E., Apprill, A., Bik, H. M., Clever, F., Connolly, S. R., De Leon, M. E., Emmett Duffy, J., Ezzat, L., Gignoux-Wolfsohn, S., Allen Herre, E., Kaye, J. Z., Kline, D. I., Kueneman, J. G., McCormick, M. K., Owen McMillan, W., O’Dea, A., Pereira, T. J., Petersen, J. M., … Eisen, J. A. (2021). Natural experiments and long-term monitoring are critical to understand and predict marine host-microbe ecology and evolution. PLoS Biology, 19(8), e3001322. 10.1371/journal.pbio.3001322

Leray, M., Yang, J. Y., Meyer, C. P., Mills, S. C., Agudelo, N., Ranwez, V., Boehm, J. T., & Machida, R. J. (2013a). A new versatile primer set targeting a short fragment of the mitochondrial COI region for metabarcoding metazoan diversity: Application for characterizing coral reef fish gut contents. Frontiers in Zoology, 10(1), 34–34.

Liedke, A. M. R., Bonaldo, R. M., Segal, B., Ferreira, C. E. L., Nunes, L. T., Burigo, A. P., Buck, S., Oliveira-Santos, L. G. R., & Floeter, S. R. (2018). Resource partitioning by two syntopic sister species of butterflyfish (Chaetodontidae). Journal of the Marine Biological Association of the United Kingdom, 98(7), 1767–1773. 10.1017/s0025315417001321

Machida, R. J., Leray, M., Ho, S.-L. L., & Knowlton, N. (2017). Metazoan mitochondrial gene sequence reference datasets for taxonomic assignment of environmental samples. Scientific Data, 4, sdata201727–sdata201727. 10.1038/sdata.2017.27

MacNally, R. C. (1995). Ecological versatility and community ecology. Cambridge University Press; Cambridge Core. 10.1017/CBO9780511565427

McMahon, K. W., Thorrold, S. R., Houghton, L. A., & Berumen, M. L. (2016). Tracing carbon flow through coral reef food webs using a compound-specific stable isotope approach. Oecologia, 180(3), 809–821. 10.1007/s00442-015-3475-3

McMurdie, P. J., & Holmes, S. (2013). phyloseq: An R Package for reproducible interactive analysis and graphics of microbiome census data. PLoS ONE, 8(4), e61217–e61217. 10.1371/journal.pone.0061217

Morais, R. A., & Bellwood, D. R. (2019). Pelagic subsidies underpin fish productivity on a degraded coral reef. Current Biology, 29(9), 1521–1527.e6. 10.1016/j.cub.2019.03.044

Morales-Nin, B. (1991). Determinación del crecimiento de peces óseos en base a la microestructura de los otolitos. FAO Documento Técnico de Pesca. https://www.fao.org/fishery/en/publications/74849

Nagelkerken, I., van der Velde, G., Wartenbergh, S. L. J., Nugues, M. M., & Pratchett, M. S. (2009). Cryptic dietary components reduce dietary overlap among sympatric butterflyfishes (Chaetodontidae). Journal of Fish Biology, 75(6), 1123–1143. 10.1111/j.1095-8649.2009.02303.x

Nielsen, J. M., Clare, E. L., Hayden, B., Brett, M. T., & Kratina, P. (2017). Diet tracing in ecology: Method comparison and selection. Methods in Ecology and Evolution. 10.1111/2041-210x.12869

Ogle, D. H., Wheeler, P., & Dinno, A. (2020). Simple Fisheries Stock Assessment Methods [R package FSA version 0.8.30].

Parravicini, V., Casey, J. M., Schiettekatte, N. M. D., Brandl, S. J., Pozas-Schacre, C., Carlot, J., Edgar, G. J., Graham, N. A. J., Harmelin-Vivien, M., Kulbicki, M., Strona, G., & Stuart-Smith, R. D. (2020). Delineating reef fish trophic guilds with global gut content data synthesis and phylogeny. PLoS Biology, 18(12 December), e3000702. 10.1371/journal.pbio.3000702

Pitts, P. A. (1991). Comparative use of food and space by three Bahamian butterflyfishes. Bulletin of Marine Science, 48(3), 749–756.

Pompanon, F., Deagle, B. E., Symondson, W. O. C., Brown, D. S., Jarman, S. N., & Taberlet, P. (2012). Who is eating what: Diet assessment using next generation sequencing. Molecular Ecology, 21(8), 1931–1950. 10.1111/j.1365-294X.2011.05403.x

Pratchett, M. S. (2005). Dietary overlap among coral-feeding butterflyfishes (Chaetodontidae) at Lizard Island, northern Great Barrier Reef. Marine Biology, 148(2), 373–382. 10.1007/s00227-005-0084-4

Pratchett, M. S., Thompson, C. A., Hoey, A. S., Cowman, P. F., & Wilson, S. K. (2018). Effects of coral bleaching and coral loss on the structure and function of reef fish assemblages. In Coral Bleaching. Ecological Studies, vol. 233, van Oppen, M. & Lough, J., eds (pp. 265–293). Springer, Cham. http://link.springer.com/10.1007/978-3-319-75393-5_11

Pratchett, M. S., Wilson, S. K., Berumen, M. L., & McCormick, M. I. (2004). Sublethal effects of coral bleaching on an obligate coral feeding butterflyfish. Coral Reefs, 23(3), 352–356.

Price, E. L., Sertić Perić, M., Romero, G. Q., & Kratina, P. (2019). Land use alters trophic redundancy and resource flow through stream food webs. Journal of Animal Ecology, 88(5), 677–689. 10.1111/1365-2656.12955

Puebla, O., Picq, S., Lesser, J. S., & Moran, B. (2018). Social-trap or mimicry? An empirical evaluation of the *Hypoplectrus unicolor*–*Chaetodon capistratus* association in Bocas del Toro, Panama. Coral Reefs, 37(4), 1127–1137. 10.1007/s00338-018-01741-0

Pulliam, H. R. (1975). Diet optimization with nutrient constraints. The American Naturalist, 109(970), 765–768. 10.1086/283041

R. Development Core Team. (2008). R: A language and environment for statistical computing [Computer software]. R Foundation for Statistical Computing.

Randall, J. E. (1967). *Food habits of reef fishes of the West Indies* (Stud. Trop). Hawaii Institute of Marine Biology. University of Hawaii, Honolulu.

Rocha, L. A., Jogan, J., Király, G., Feráková, V., & Bernhardt, K.-G. (2010). *Chaetodon capistratus*. https://www.iucnredlist.org/species/165695/6094300

Rodas, A. M., Wright, R. M., Buie, L. K., Aichelman, H. E., Castillo, K. D., & Davies, S. W. (2020). Eukaryotic plankton communities across reef environments in Bocas del Toro Archipelago, Panamá. Coral Reefs, 1–15. 10.1007/s00338-020-01979-7

Roehr, J. T., Dieterich, C., & Reinert, K. (2017). Flexbar 3.0 – SIMD and multicore parallelization. Bioinformatics, 33(18), 2941–2942. 10.1093/bioinformatics/btx330

Rognes, T., Flouri, T., Nichols, B., Quince, C., & Mahé, F. (2016). VSEARCH: A versatile open source tool for metagenomics. PeerJ, 2016(10). 10.7717/peerj.2584

Rotjan, R. D., & Lewis, S. M. (2008). Impact of coral predators on tropical reefs. Marine Ecology Progress Series, 367, 73–91. 10.3354/meps07531

Seemann, J., Yingst, A., Stuart-Smith, R. D., Edgar, G. J., & Altieri, A. H. (2018). The importance of sponges and mangroves in supporting fish communities on degraded coral reefs in Caribbean Panama. PeerJ, 2018(3), e4455–e4455. 10.7717/peerj.4455

Semmler R. F., Brandl S. J., Keith S. A., Bellwood D. R. (2021). Fine-scale foraging behavior reveals differences in the functional roles of herbivorous reef fishes. Ecology Evolution, 11: 4898–4908. 10.1002/ece3.7398

Semmler, R. F., Sanders, N. J., CaraDonna, P. J., Baird, A. H., Jing, X., Robinson, J. P. W., Graham, N. A. J., & Keith, S. A. (2022). Reef fishes weaken dietary preferences after coral mortality, altering resource overlap. Journal of Animal Ecology, 91, 2125–2134. 10.1111/1365-2656.13796

Tesch, F. W. (1968). Age And Growth, Methods for assessment of fish production in fresh waters (W. E. Ricker, Ed.). Blackwell Scientific.

Traugott, M., Thalinger, B., Wallinger, C., & Sint, D. (2020). Fish as predators and prey: DNA -based assessment of their role in food webs. Journal of Fish Biology, jfb.14400-jfb.14400. 10.1111/jfb.14400

Uchida, S., Drossel, B., & Brose, U. (2007). The structure of food webs with adaptive behaviour. Ecological Modelling, 206(3–4), 263–276. 10.1016/j.ecolmodel.2007.03.035

Van Leeuwen, E., Brannstrom, A., Jansen, V. A. A., Dieckmann, U., & Rossberg, A. G. (2013). A generalized functional response for predators that switch between multiple prey species. Journal of Theoretical Biology, 328, 89–98. 10.1016/j.jtbi.2013.02.003

Venables, W. N., & Ripley, B. D. (2002). *Modern Applied Statistics with S* (4th ed). Springer. https://www.stats.ox.ac.uk/pub/MASS4/.

Vestheim, H., & Jarman, S. N. (2008). Blocking primers to enhance PCR amplification of rare sequences in mixed samples—A case study on prey DNA in Antarctic krill stomachs. Frontier in Zoology, 5, 12–12. 10.1186/1742-9994-5-12

Vrolijk, N. H., Targett, N. M., Woodin, B. R., & Stegeman, J. J. (1995). Comparison of cytochrome P450 in three butterflyfish species: Ecological implications of elevated CYP2B and CYP3A in *Chaetodon capistratus*. Marine Environmental Research, 39(1), 11–14. 10.1016/0141-1136(94)00076-2

Whiteman, E. A., Côté, I. M., & Reynolds, J. D. (2007). Ecological differences between hamlet (*Hypoplectrus*: Serranidae) colour morphs: Between-morph variation in diet. Journal of Fish Biology, 71(1), 235–244. 10.1111/j.1095-8649.2007.01485.x

Wilson, S.K., Graham, N.A.J., Pratchett, M.S., Jones, G.P. and Polunin, N.V.C. (2006), Multiple disturbances and the global degradation of coral reefs: are reef fishes at risk or resilient?. Global Change Biology, 12: 2220–2234. 10.1111/j.1365-2486.2006.01252.x

Wong, B. B. M., & Candolin, U. (2015). Behavioral responses to changing environments. Behavioral Ecology, 26(3), 665–673. 10.1093/beheco/aru183

