## Supplementary File for "Dietary resilience of coral reef fishes to habitat degradation"

Contents

Supplementary Figures	2

Supplementary Tables	14

Supplementary Methods	16

References	17

**Supplementary figures**

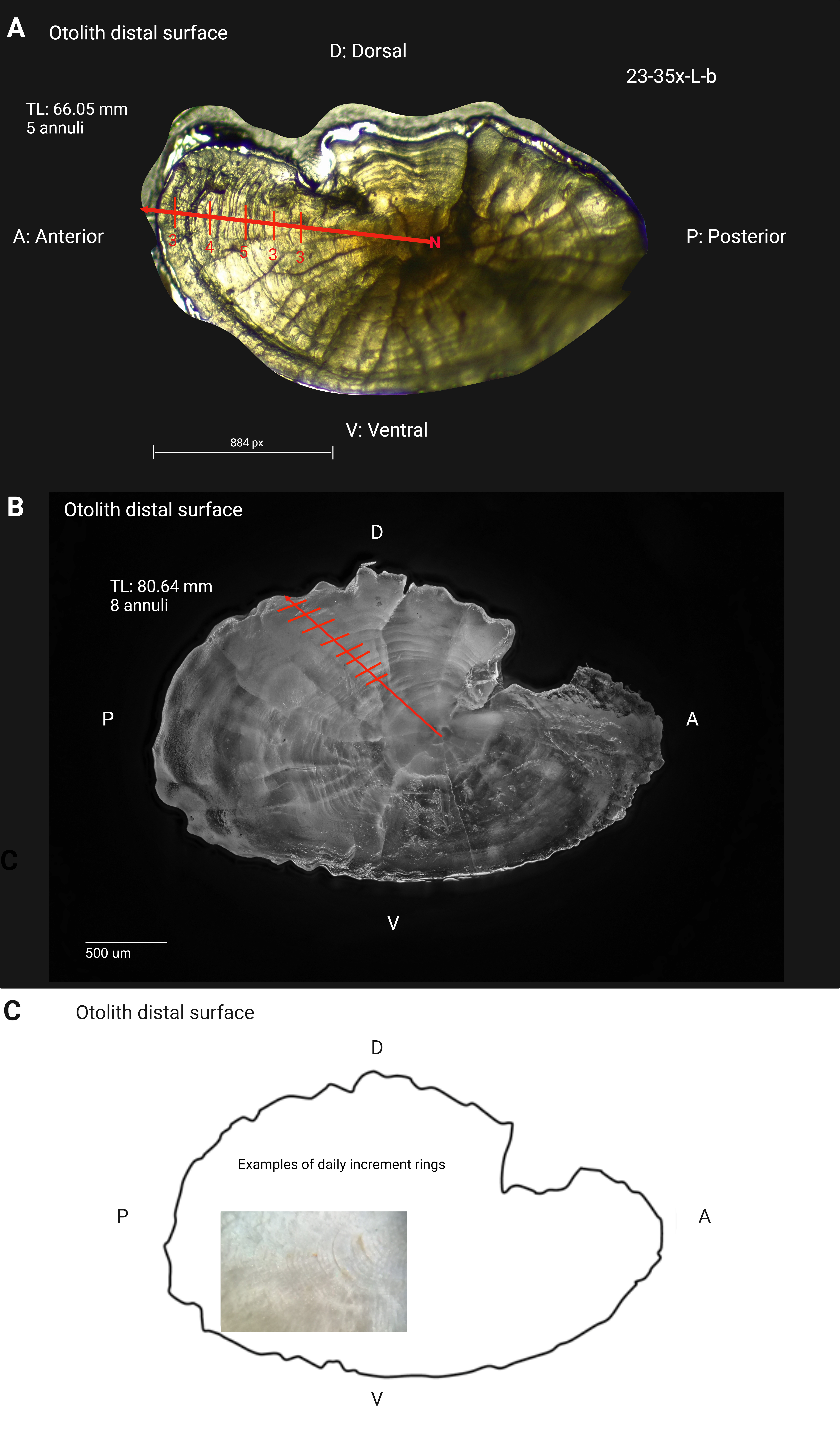

**Fig. S1.** Depicted are the distal surfaces of the (A) left and (B) right sagittal otolith of a *Chaetodon capistratus* fish individual; (C) right sagittal otolith with visible daily increments.

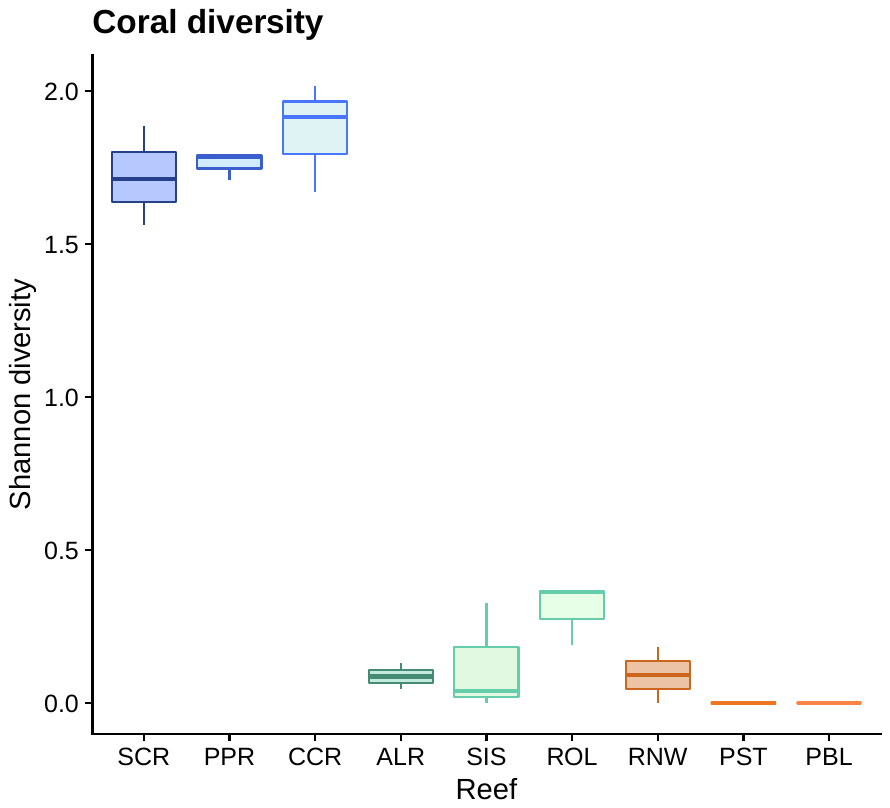

**Fig. S2.** Estimates of coral diversity at nine non-contiguous reefs and across three different reef zones (blue = outer bay, green = inner bay, orange = inner bay disturbed) at the Bahía Almirante, Bocas del Toro, Panamá.

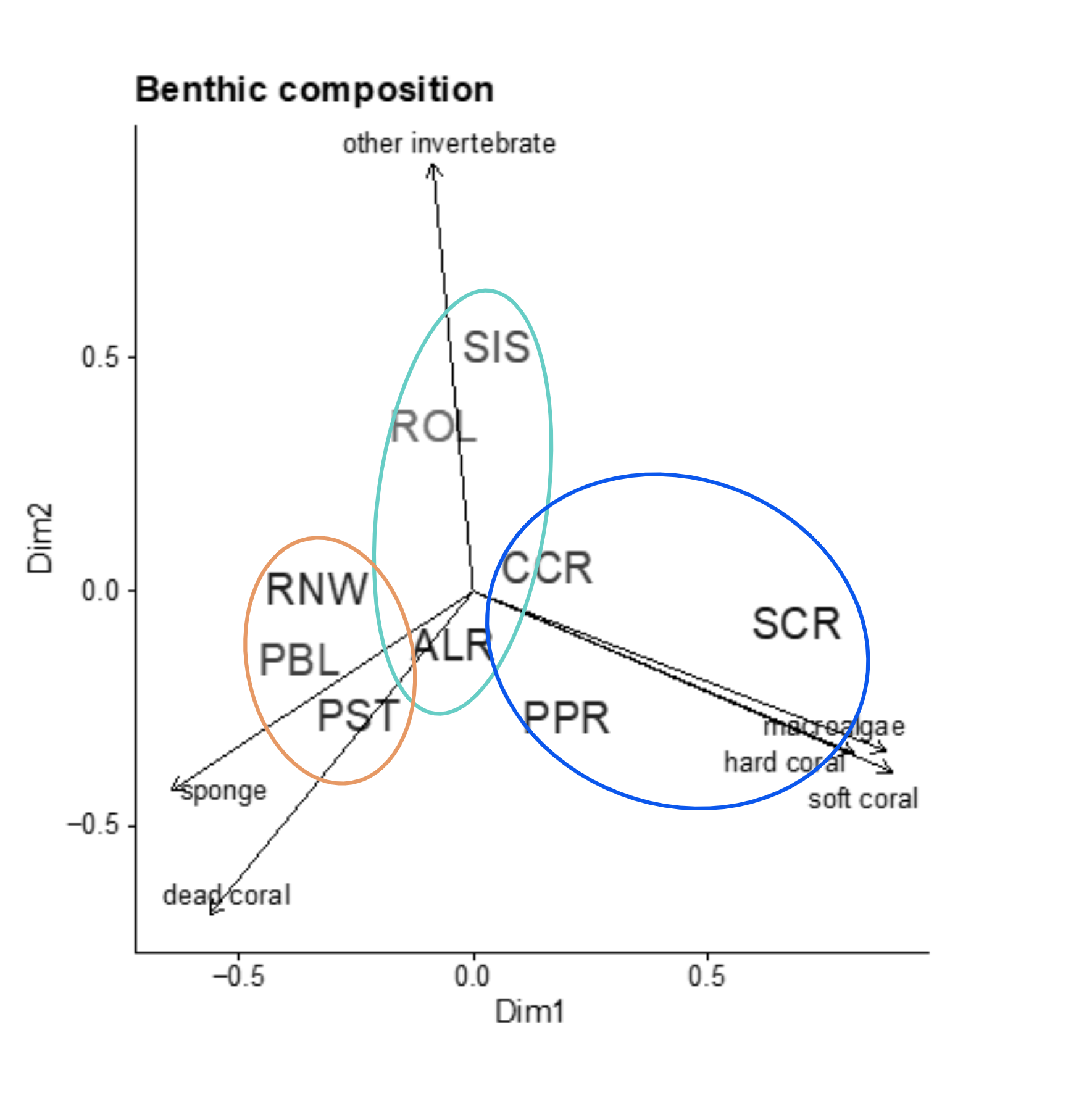

**Fig. S3.** Principal coordinates analysis (PCoA) of benthic composition at nine study reefs across three reef zones (blue = outer bay, green = inner bay, orange = inner bay disturbed) at the Bahìa Almirante, Bocas del Toro, Panamá. Vectors overlayed indicate the major benthic groups or substrate types responsible for driving differences between the nine study reefs.

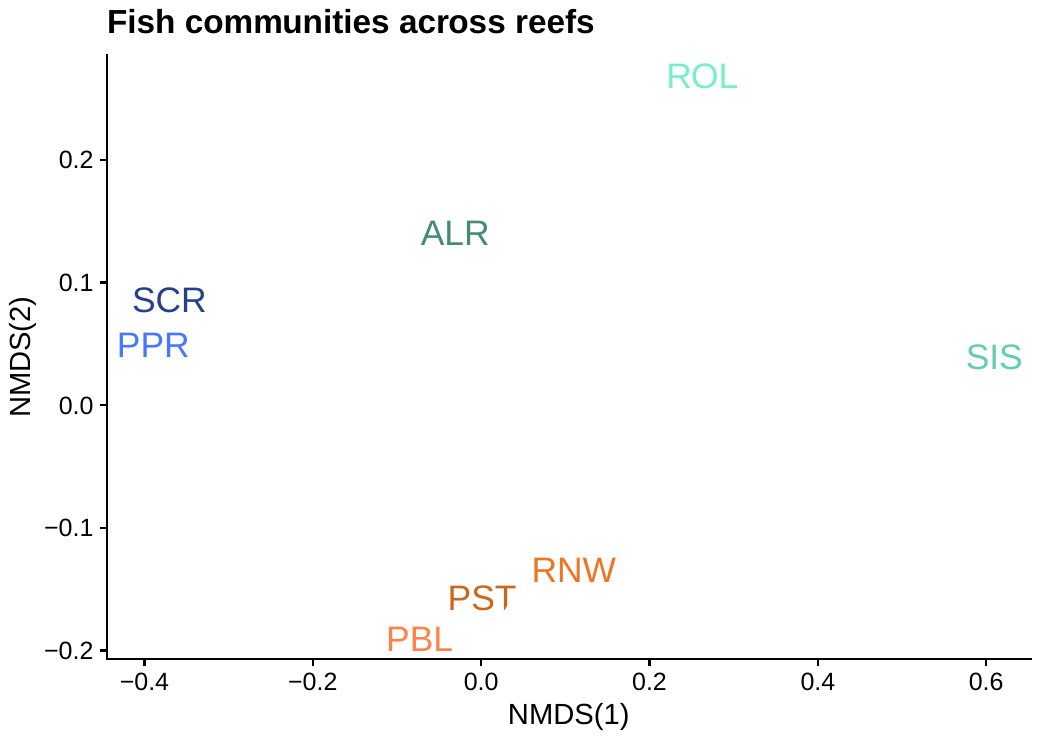

**Fig. S4.** Nonmetric multidimensional scaling (NMDS) using Bray Curtis dissimilarity of fish communities across reefs and zones (color coded): blue = outer bay, green = inner bay, red = inner bay disturbed. Fish surveys were conducted at all of our nine study reefs except of Coral Caye Reef (CCR) in the outer bay zone.

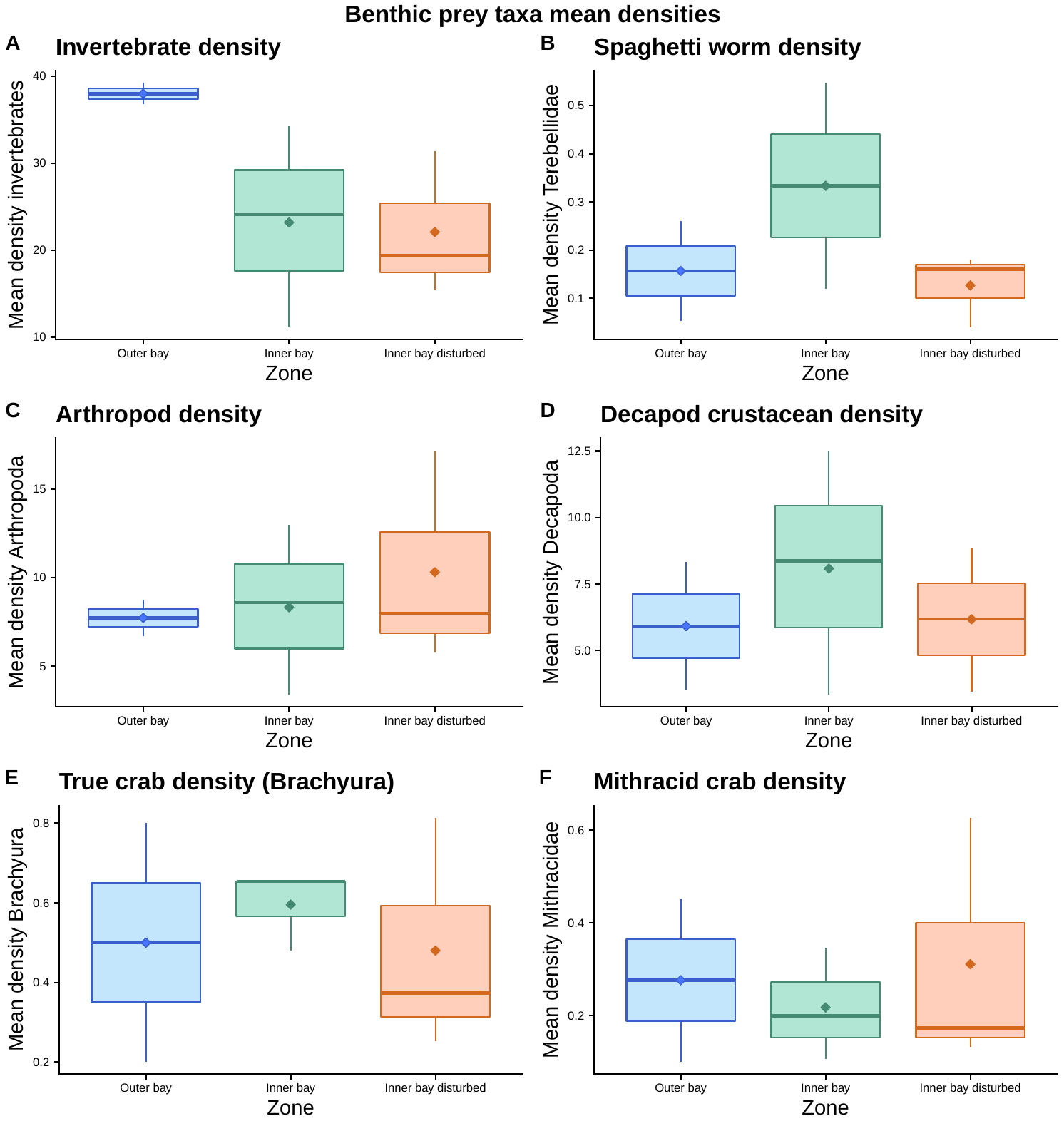

**Fig. S5.** Invertebrate mean densities representing potential prey taxa of two focal fish species at our study area across three reef zones from highest coral cover (outer bay) to lowest coral cover (inner bay disturbed). Depicted are mean densities of (A) all recorded benthic macro-invertebrates (> 2 mm); (B) terebellid worms, a main diet item of *C. capistratus*; (C) Arthropods, the main prey of *H. puella*. In addition, we examined lower taxonomic levels within the phylum Arthropoda to examine potential differences among zones that might influence fish diet: (D) decapod crustaceans; (E) brachyuran crabs and (F) mithracid crabs. Macro-invertebrates (> 2 mm) were collected within three quadrats per reef (50 x 50 cm) and on three reefs per zone.

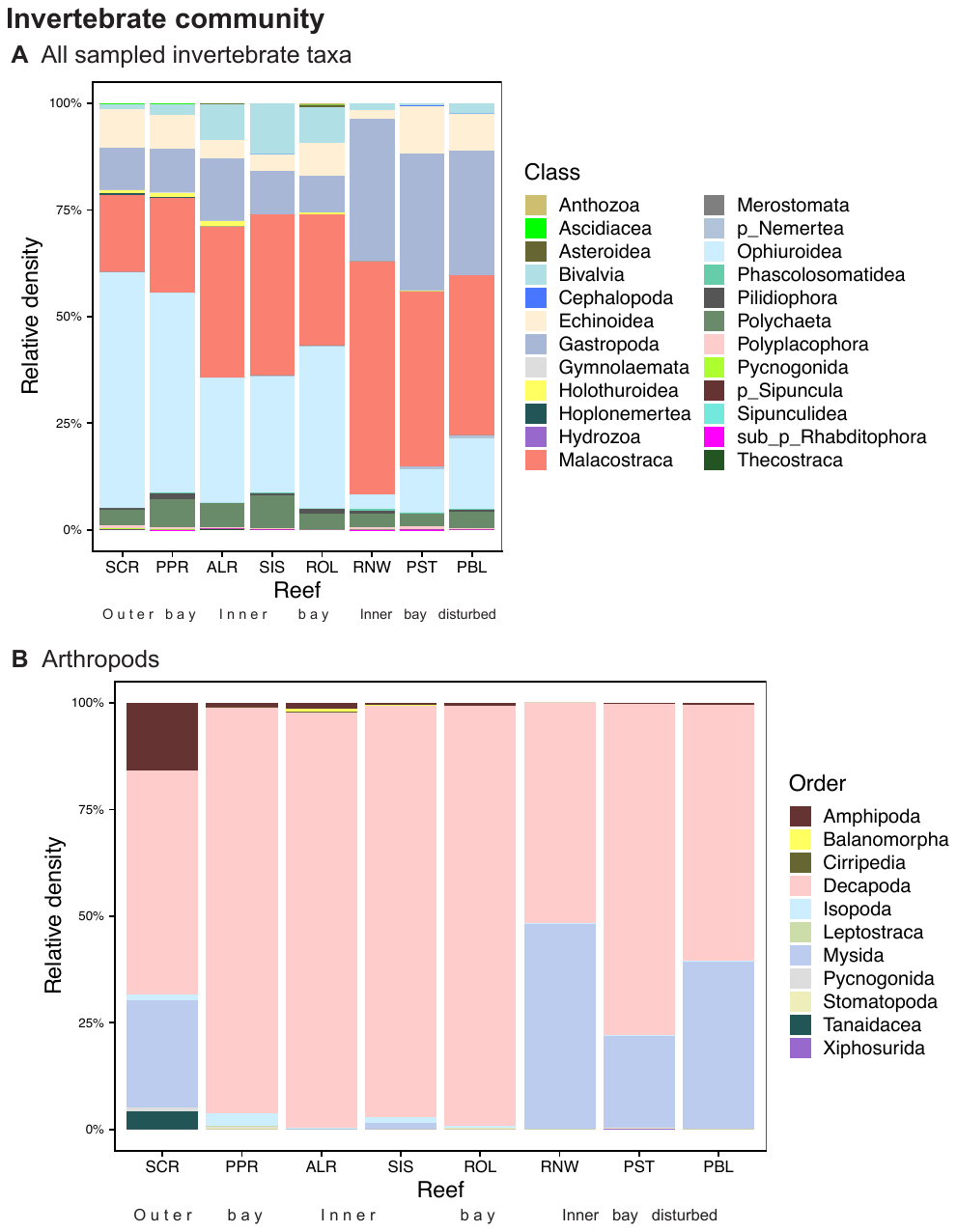

**Figure S6.** (A) Invertebrate relative densities collected using quadrats on dead patches (predominantly *Agaricia tenuifolia*) at eight study reefs depicted at the taxonomic level of class; where the level of class was not identified, the next higher taxonomic level identified is depicted indicated by p = phylum or sub_p = subphylum. (B) the relative densities of arthropod taxa collected from the same quadrats as (A).

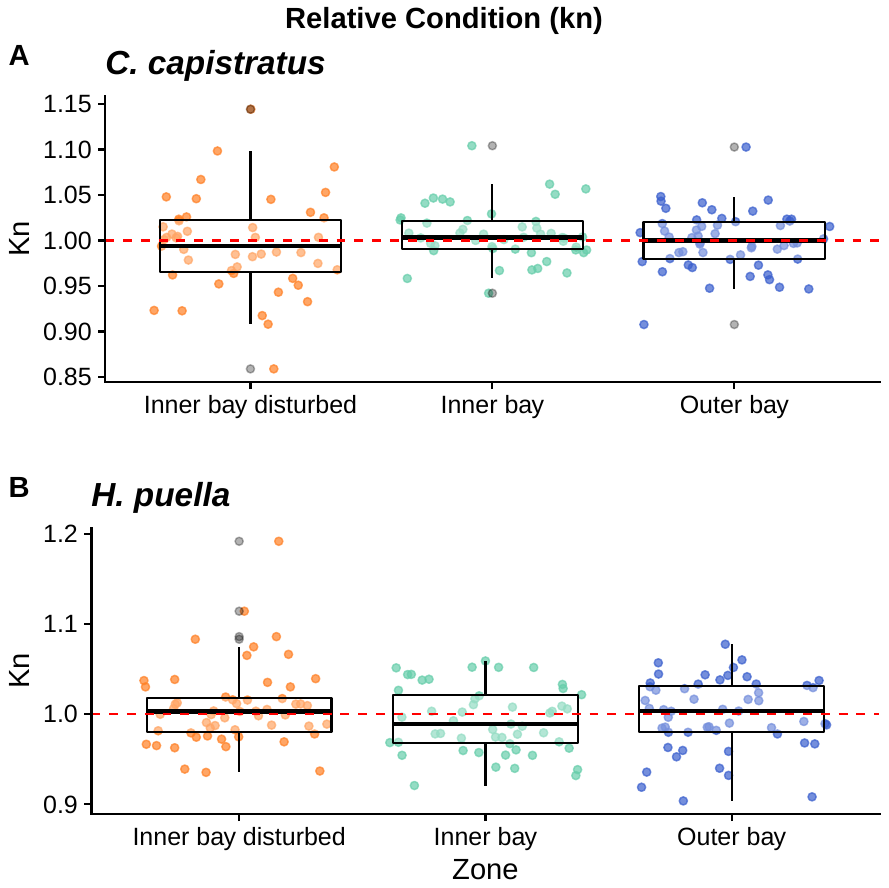

**Fig. S7.** Relative condition factor (kn) by zone not taking into account size classes for (A) *Chaetodon capistratus* and (B) *Hypoplectrus puella*. Values of 1.0 (red line) and above represent optimal condition, whereas values below 1.0 indicate suboptimal condition.

A

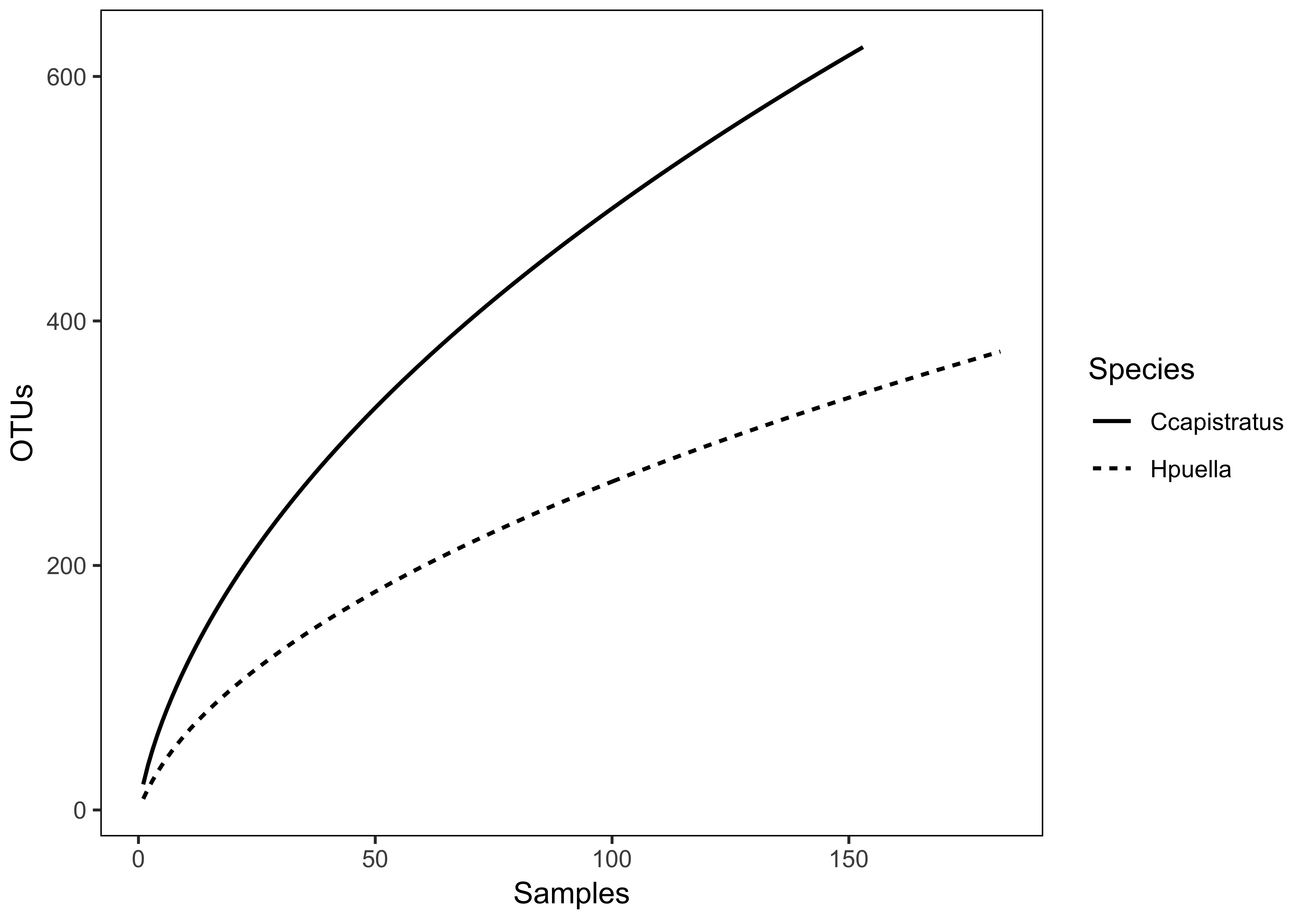

B	C

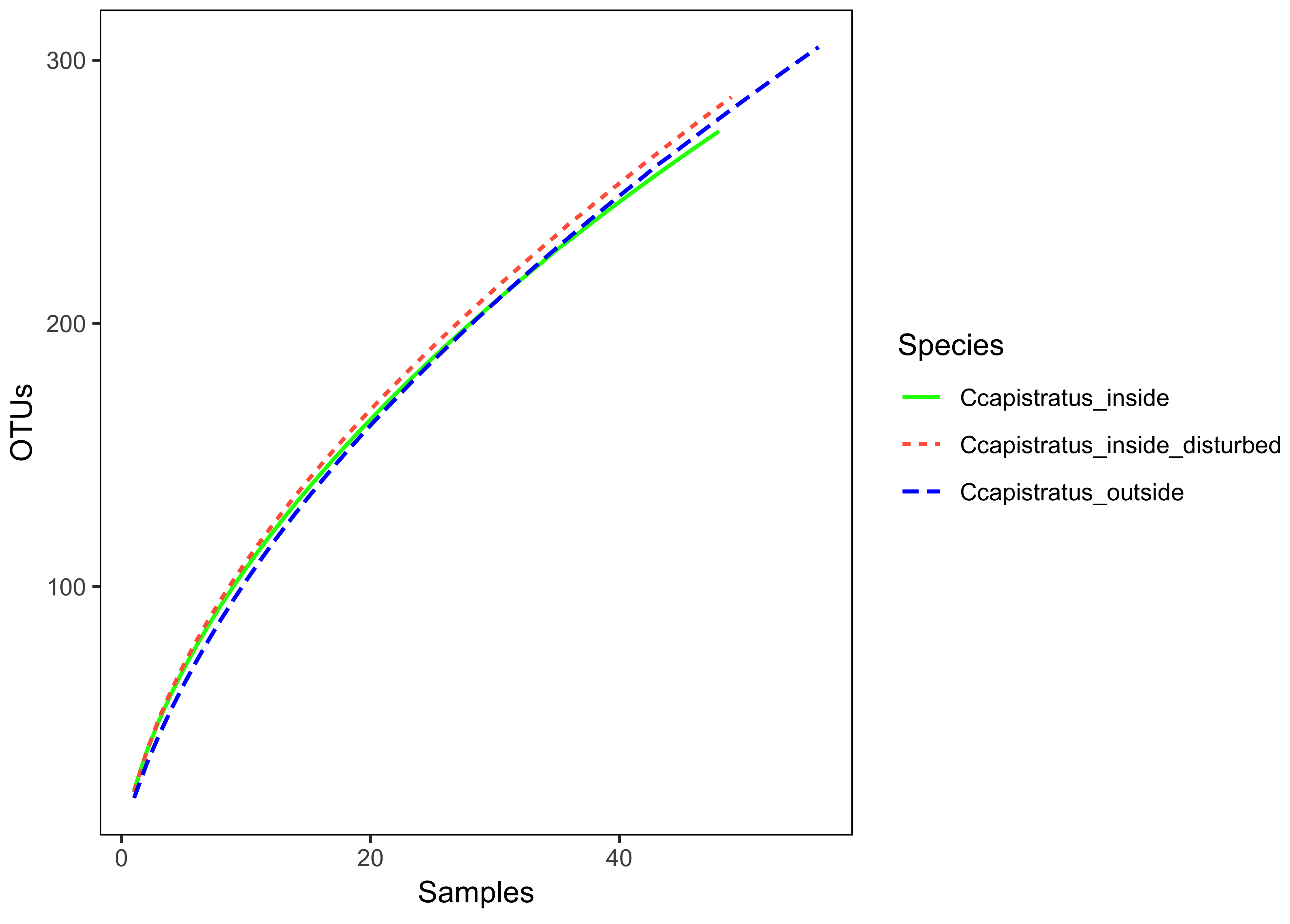

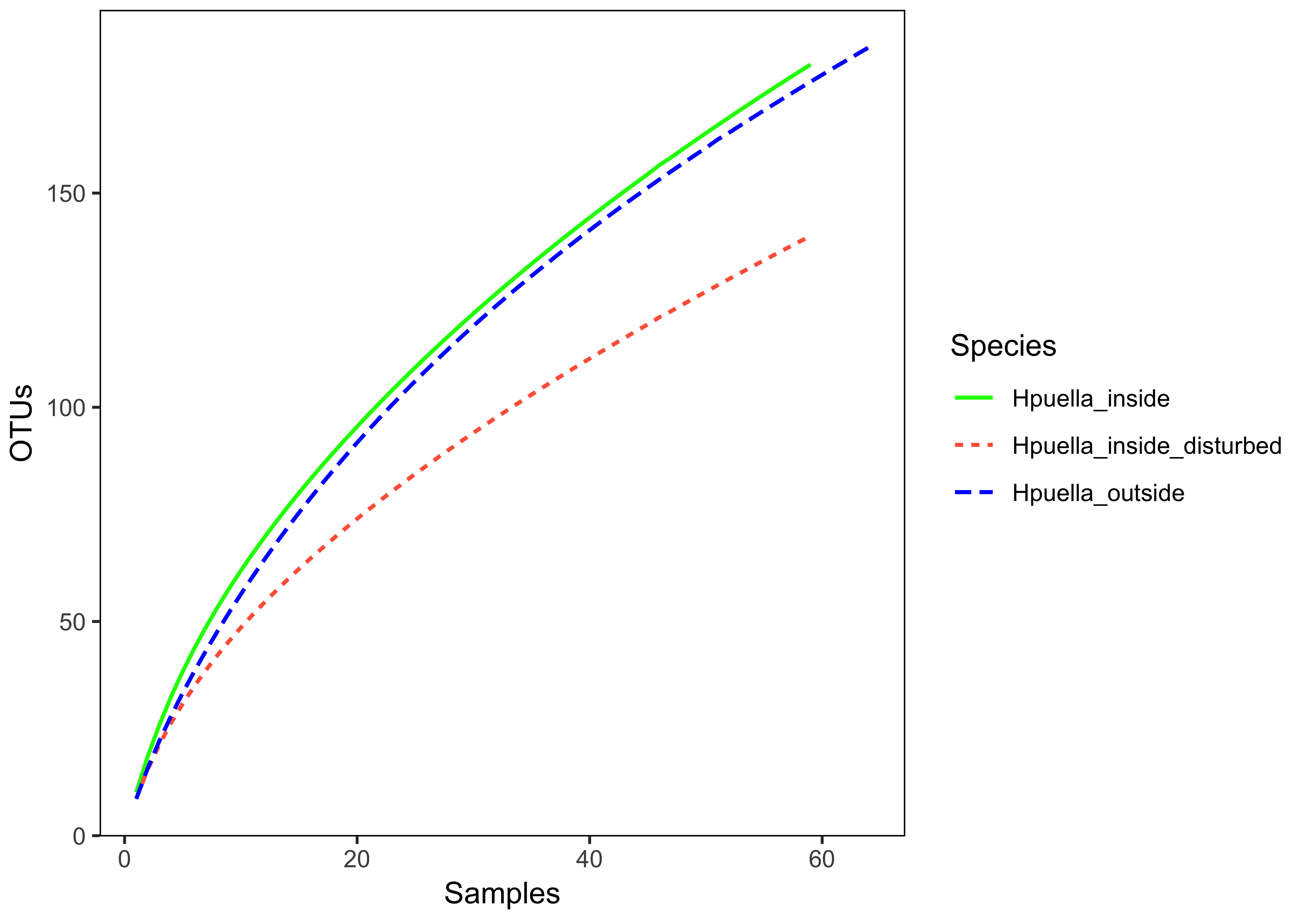

Fig. S8 Sample-based rarefaction curves for (A) both study species across all samples, (B) *Chaetodon capistratus* and (C) *Hypoplectrus puella* both by reef zone.

**Distribution of sample sequencing depth**

**A *C. capistratus***

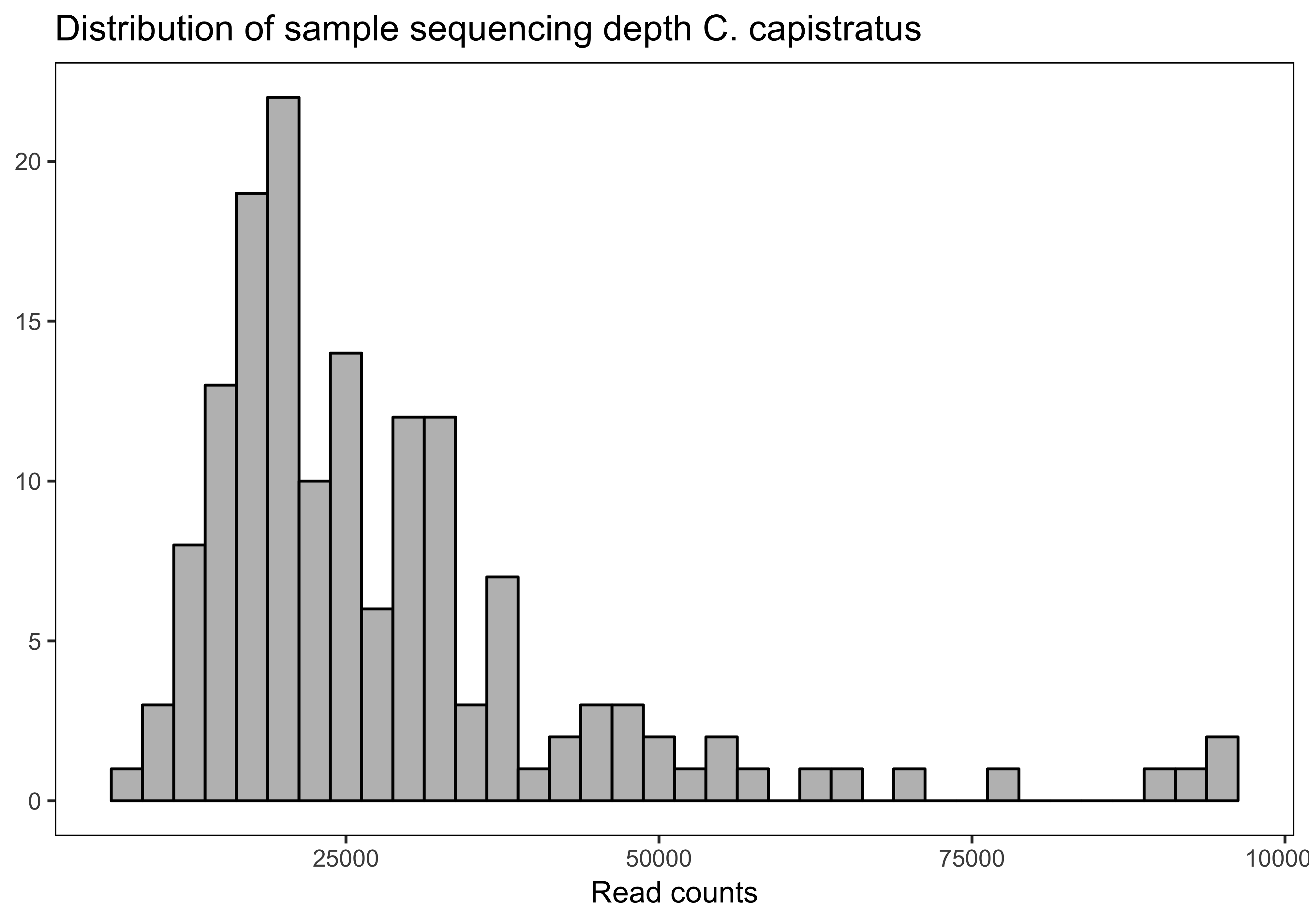

**B *H. puella***

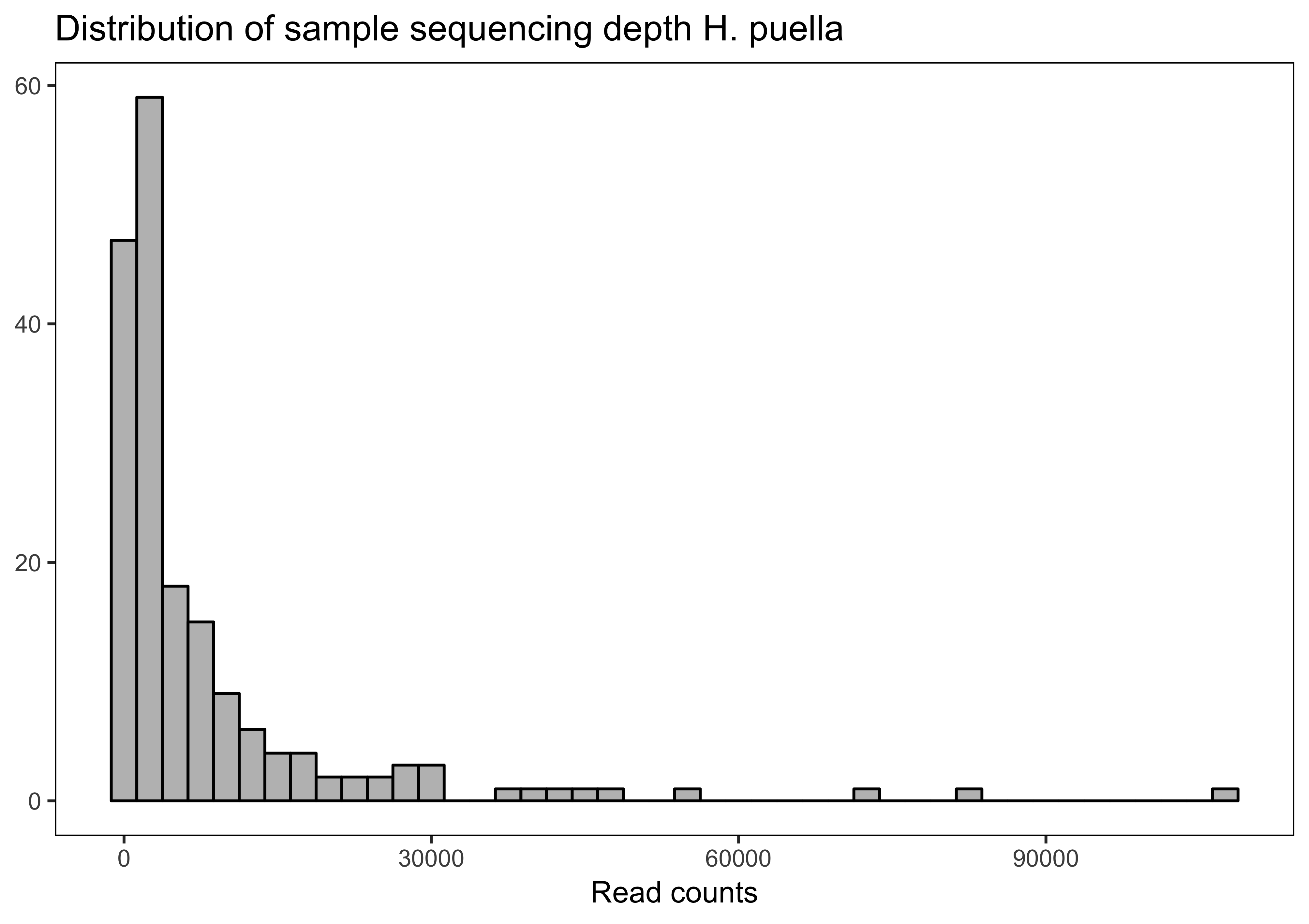

Fig. S9 The distribution of sequencing depth (sequencing read counts) across (A) stomach content samples of *Chaetodon capistratus* and (B) intestinal content samples of *Hypoplectrus puella*.

Fig. S9 The distribution of sequencing depth (sequencing read counts) across (A) stomach content samples of *Chaetodon capistratus* and (B) intestinal content samples of *Hypoplectrus puella*.

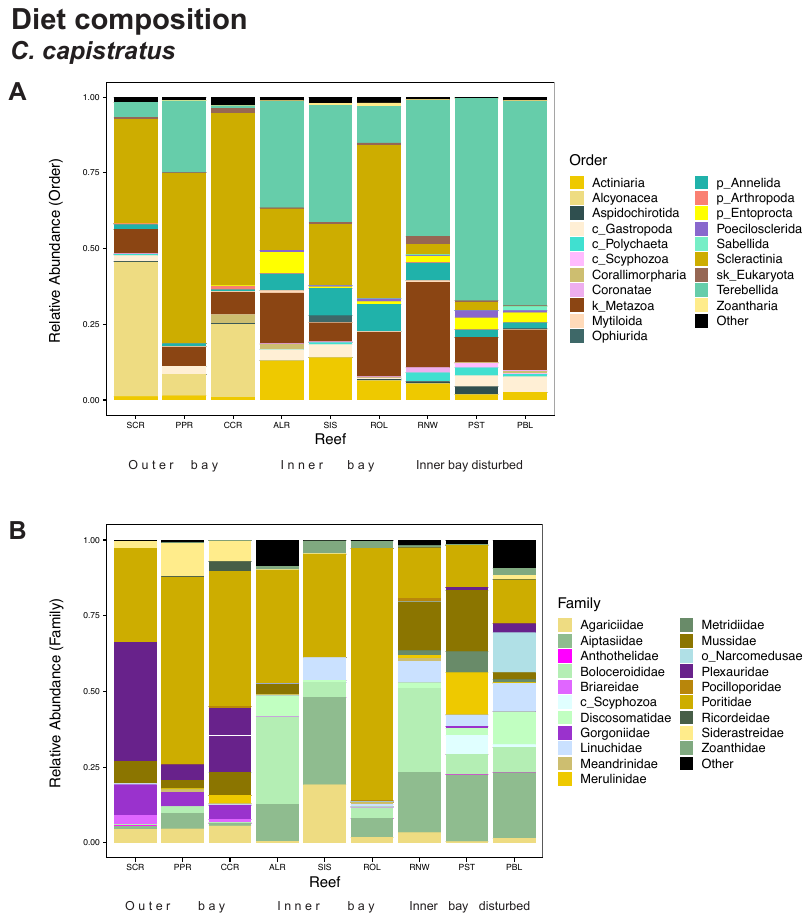

Fig. S10. *Chaetodon capistratus* diet composition across nine study reefs (A) by order for all eukaryotes found in the diet, and (B) by family within the phylum Cnidaria (or alternatively the next higher level that could be taxonomically assigned). The diet changes along the coral cover gradient from Salt Creek (SCR, highest coral cover, far left) to Punta Puebla (PBL, lowest coral cover, far right).

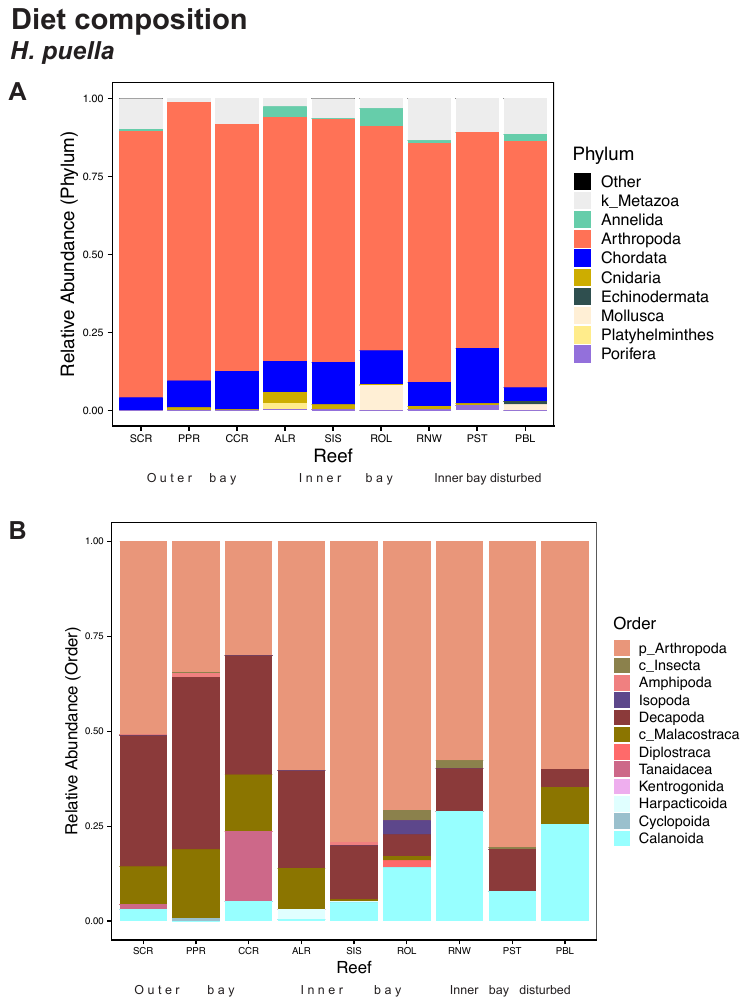

Fig. S11 (A): *Hypoplectrus puella* diet composition showing the nine most common prey phyla (or alternatively the next higher level that could be taxonomically assigned (i.e., k = kingdom) across nine study reefs. For the carnivore *H. puella*, we only included OTUs that were taxonomically assigned to kingdom Metazoa. Arthropods dominate the diet across all reefs and habitat zones despite different levels of coral cover (i.e., from Salt Creek (SCR, highest coral cover, far left) to Punta Puebla (PBL, lowest coral cover, far right). (B): Within only arthropods at the order level (or alternatively the next higher level that could be taxonomically assigned; i.e., c = class, p = phylum), it becomes apparent that more planktonic taxa (i.e., calanoid copepods) are consumed with decreasing coral cover (towards the right) and more decapods (including class Malacostraca) with increasing coral cover (towards the left).

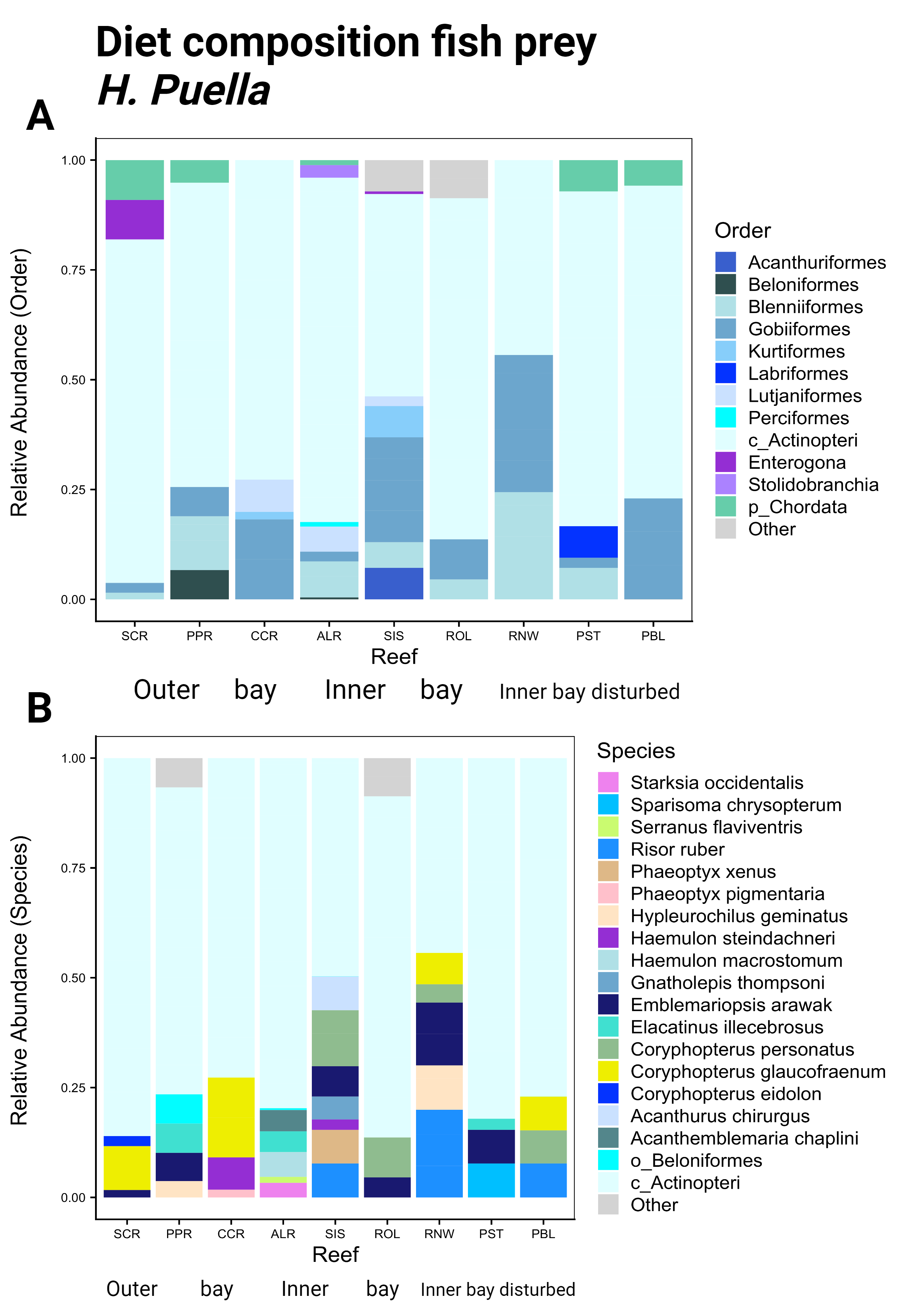

Fig. S12 *Hypoplectrus puella* diet composition across nine study reefs (A) within phylum Chordata by order (or alternatively the next higher level that could be taxonomically assigned i.e., c = class; p = phylum) and (B) including only fishes at the species level (or alternatively the next higher level that could be taxonomically assigned i.e., o = order; c = class).

**Supplementary Tables**

**Table S1.** The versatile COI primer pair (Leray 2013) that was used in this study

| **Primer_name** | **Primer_sequence (5'-3')** |
| --- | --- |
| mlCOIintF | GGWACWGGWTGAACWGTWTAYCCYCC |
| jgHCO2198 | TAIACYTCIGGRTGICCRAARAAYCA |

**Table S2.** Species-specific blocking primer sequences for two coral reef fishes, *Hypoplectrus puella* and *Chaetodon capistratus.*

| **Primer_name** | **Primer_sequence (5'-3')** |
| --- | --- |
| Hpuella-Blocker | CAAAGAATCAGAATAGATGTTGGTAAAGA-C3 |
| Ccapistratus-Blocker | CAAAGAATCAGAACAGGTGTTGGTAAAGA-C3 |

**Table S3.** One-way Analysis of Variance (ANOVA) results testing whether the mean densities of invertebrates overall, specific groups and taxa differed significantly among three reef zones comprising two (outer bay) or three (inner bay, inner bay disturbed) reefs (n = 8). Invertebrates (> 2 mm) were collected from dead coral habitat using three quadrates (50 x 50 cm) per reef. Significant differences are depicted in bold.

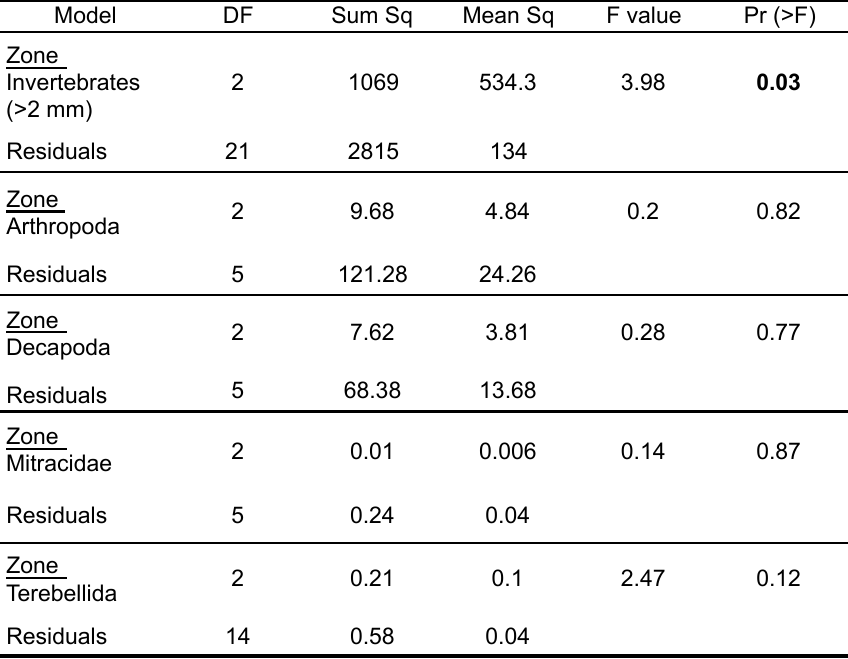

**Table S4.** Post hoc Tukey test results showing which pairwise comparisons of invertebrate mean densities are responsible for significant differences among three zones as detected by ANOVA (Table S3). Invertebrate mean densities significantly differed between the outer bay zone and inner bay disturbed zone (depicted in bold).

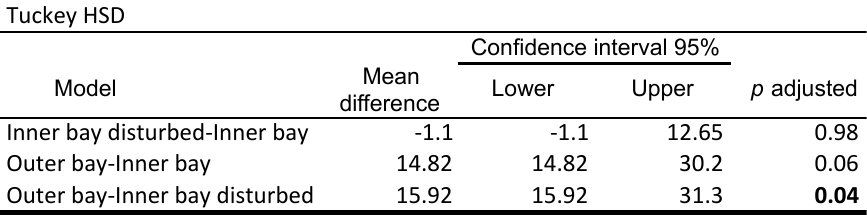

**Supplementary Methods**

**I. Study system**
The bay is confined by the mainland and protected from ocean swell by several islands leading to restricted water-exchange with the open ocean. Together with local climatic conditions, this creates an environment with limited water flow, variable salinity (fluctuating locally from 20–34 PSS) (Kaufmann and Thompson 2005; Collin et al. 2009) and elevated sea surface temperatures during calm weather periods (Cramer 2013; Altieri et al. 2017). While terrestrial run-off naturally elevates nutrients within the bay, both untreated wastewater from tourism development and agricultural discharges intensify eutrophic levels (Guzmán et al. 2005; D’Croz, Rosario, and Gondola 2005; Cramer 2013; Altieri et al. 2017).

**II. Benthic, fish and invertebrate surveys**
Three replicate transect lines (20 m) per reef were placed parallel to the shore at a depth of 2-4 m. To estimate benthic cover and community composition, ten quadrats (100 x 70 cm) were photographed at two meter increments along each transect (quadrats per site N = 30). We analyzed each photo using CoralNet (Beijbom et al. 2015) with a grid of 100 points over each photo, and the identity of the primary space holder under each point was determined to the level of broad taxonomic groups (e.g., hard coral, soft coral, macroalgae, sponge, dead coral, zoanthids, rubble) to avoid potential identification errors arising from variation in image quality. Hard corals were identified at genus level where possible. Fish communities were surveyed along a 20 x 5 m belt by one experienced surveyor recording the abundance and identity of all non-cryptic fish species using scuba. To assess macro-invertebrate (> 2 mm) diversity and abundance, quadrats (50 x 50 cm) were placed on continuous surfaces of dead coral (mainly *Agaricia tenuifolia*) and all coral rubble was carefully collected in bins and transported to the field station. At the laboratory, all invertebrates larger than 2mm were counted and identified to the lowest taxonomic level.

**III. Fish collection**
To avoid cross-contamination between fish samples*,* we strictly followed the following procedures during fish sample preparation and DNA extraction: First, sterile gloves were changed. Second, the flow-hood was DNA de-contaminated using 10% Sodium Hypochlorite followed by a 90% EtOH rinse. Third, scissors and forceps were placed in a 10% Sodium Hypochlorite bath for 15 min immediately after use, thoroughly rinsed with Milli-Q water and subsequently 70% ethanol and flamed to remove any remaining bleach, water contaminants or tissue.

**IV.** **DNA Extraction**
Between 0.05 and 0.25 g of prey tissue per sample from *C. capistratus*’ stomach contents and between 0.05 and 0.25 g of *H. puella* digesta (intestinal contents) were added to individual Eppendorf tubes containing power beads with bead solution, C1 solution (60 ul) and 20 ul proteinaseK (0.4 mg.mL-1) to enhance the lysis of animal tissue as well as algae. Because yield was low in an initial set of five extractions of intestinal content samples, we briefly vortexed the Eppendorf tubes with digesta before an additional incubation step of 15 min at 60°*C* with 1000 rpm agitation to allow for beginning of tissue lysis by Proteinase K before mechanical disruption by vortexing with beads. Samples were then vortexed for 5 min on the Vortex Genie 2 with vortex adapter followed by a 105 min incubation at 60°*C* with 1000 rpm agitation. The Eppendorf tubes containing stomach content samples were vortexed for 5 min using a Vortex Genie 2 (Scientific Industries) with vortex adapter and subsequently incubated at 60°*C* for two hours with agitation (1000 rpm) on an Eppendorf™ Thermomixer™ R. The longer incubation step recommended in previous studies (Leray, Meyer, and Mills 2015; Wangensteen and Turon 2017) helped lyse hard-shelled invertebrates as well as coral and potentially algae. DNA extracts were diluted 5 times in nuclease-free water and the molecular weight of the extracted genomic DNA was assessed with electrophoresis of GelRed™-stained DNA on a 1.5% agarose gel. DNA concentration (ng/ul) was quantified with a Quant-iT™ dsDNA High-Sensitivity Assay Kit using an Invitrogen Qubit® Fluorometer (Life Technologies), before storing DNA extracts at *−*20*^◦^C*.

**V.** **Metabarcoding library preparation**
We followed a previously published protocol for sample multiplexing that uses a combination of tailed PCR primers and ligation of indexed adapters (Leray, Haenel, and Bourlat 2016). To do so we used matching oligonucleotide indices (Binladen et al. 2007; Coissac, Riaz, and Puillandre 2012) on both forward and reverse primers to prevent tag-jumping—a process that may generate spurious assignments of sequence reads to samples (Schnell, Bohmann, and Gilbert 2015). A total volume of 20 ul was used in each PCR reaction comprised of 2 x PCR buffer (Clonetech) with 1.8 mM MgCl2, 3% DMSO, 0.2 mM dNTP, 0.4 Advantage TAQ polymerase (Clonetech), 1 M of forward and reverse primer respectively (mlCOIintF and jgHCO2198) (Leray et al. 2013; Geller et al. 2013) and 1 ng/ul of DNA template. A PCR blank (using 1 ng/l nuclease free water instead of DNA template) was included in each PCR replicate run, and positive controls were included in PCRs of gut samples. Each PCR thermal cycle consisted of an initial denaturation step of 5 minutes at 95*^◦^C* followed by 38 cycles of 95*^◦^C* (30 seconds), 48*^◦^C* (30 seconds), 72*^◦^C* (45 seconds), with a final 5 minute extension at 72*^◦^C* and a final cooling step of 4*^◦^C*. PCRs were assessed with electrophoresis on 1.5% agarose gel stained with GelRed™. The three PCR replicates generated for each sample were pooled. PCR product clean-ups were performed using DNA Purification Solid Phase Reversible Immobilization (SPRI) magnetic beads (KAPA Pure Beads, Roche) using a bead:DNA ratio of 1.6:1. To achieve similar numbers of reads per sample after sequencing, cleaned-up amplicon DNA was normalized at 5ng/ul using nuclease-free water. Equimolar amplicon DNA of samples with each unique tag were then pooled into the respective adapter groups for adapter ligation. The TruSeq DNA PCR-free LT library Prep Kit (Illumina) was used for library preparation following the manufacturer's protocol.

**VI. Generalized linear mixed effects models**

According to our study design, we included the random effect of zone with reef nested within zone. The distribution of data was checked using histograms and transformed to optimize models i.e., hard coral was square root-, annelid fourth root-, and crustaceans log transformed. Akaike information criterion (AICc) (aictab function, AICcmodavg v2.2-2, Mazerolle 2019) was used to pre-select models. Final selection was based on likelihood ratio tests (function anova, lme4 package v1.1-21, Bates et al. 2015) testing the significance of the predictor variables against null-models. Data were visualized using q plots and model fit was examined with Q-Q plots.

**References**

Altieri, Andrew H, Seamus B Harrison, Janina Seemann, Rachel Collin, Robert J Diaz, and Nancy Knowlton. 2017. “Tropical Dead Zones and Mass Mortalities on Coral Reefs.” *Proceedings of the National Academy of Sciences of the United States of America* 114 (14): 3660–65.

Bates, Douglas, Martin Mächler, Benjamin M. Bolker, and Steven C. Walker. 2015. “Fitting Linear Mixed-Effects Models Using Lme4.” *Journal of Statistical Software* 67 (1).

Beijbom, Oscar, Peter J. Edmunds, Chris Roelfsema, Jennifer Smith, David I. Kline, Benjamin P. Neal, Matthew J. Dunlap, et al. 2015. “Towards Automated Annotation of Benthic Survey Images: Variability of Human Experts and Operational Modes of Automation.”. *PLOS ONE* 10 (7): e0130312.

Binladen, Jonas, M. Thomas P. Gilbert, Jonathan P. Bollback, Frank Panitz, Christian Bendixen, Rasmus Nielsen, and Eske Willerslev. 2007. “The Use of Coded PCR Primers Enables High-Throughput Sequencing of Multiple Homolog Amplification Products by 454 Parallel Sequencing.” *PLoS ONE* 2 (2).

Coissac, Eric, Tiayyba Riaz, and Nicolas Puillandre. 2012. “Bioinformatic Challenges for DNA Metabarcoding of Plants and Animals.” *Molecular Ecology* 21 (8): 1834–47.

Collin, Rachel, Luis D’Croz, Plinio Gondola, and Juan B. Del Rosario. 2009. “Climate and Hydrological Factors Affecting Variation in Chlorophyll Concentration and Water Clarity in the Bahia Almirante, Panama.” in *Proceedings of the Smithsonian Marine Science Symposium*, edited by Lang, Michael A., MacIntyre, Ian G., and Ruetzler, Klaus., First ed. 323–334. Smithsonian Contributions to the Marine Sciences. Washington DC: Smithsonian Institution Scholarly Press. https://doi.org/10.5479/10088/19175

Cramer, Katie L. 2013. “History of Human Occupation and Environmental Change in Western and Central Caribbean Panama.” *Bulletin of Marine Science* 89 (4): 955–82.

D’Croz, Luis, Juan B. del Rosario, and Plinio Gondola. 2005. “The Effect of Fresh Water Runoff on the Distribution of Dissolved Inorganic Nutrients and Plankton in the Bocas Del Toro Archipelago, Caribbean Panamá.” *Caribbean Journal of Science* 41 (3): 414–429.

Geller, J, C Meyer, M Parker, and H Hawk. 2013. “Redesign of PCR Primers for Mitochondrial Cytochrome c Oxidase Subunit I for Marine Invertebrates and Application in All-Taxa Biotic Surveys.” *Molecular Ecology Resources* 13 (5): 851–61.

Guzmán, Héctor M., Penelope A. G. Barnes, Catherine E. Lovelock, and Ilka C Feller. 2005. “A Site Description of the CARICOMP Mangrove, Seagrass and Coral Reef Sites in Bocas Del Toro, Panamá.” *Caribbean Journal of Science* 41 (3): 430–440.

Kaufmann, Karl W., and Ricardo C. Thompson. 2005. “Water Temperature Variation and the Meteorological and Hydrographic Environment of Bocas Del Toro, Panama.”

Leray, Matthieu, Quiterie Haenel, and Sarah J. Bourlat. 2016. “Preparation of Amplicon Libraries for Metabarcoding of Marine Eukaryotes Using Illumina MiSeq: The Adapter Ligation Method.” In *Methods in Molecular Biology*, 1452:209–18. Humana Press Inc.

Leray, Matthieu, Christopher P Meyer, and Suzanne C Mills. 2015. “Metabarcoding Dietary Analysis of Coral Dwelling Predatory Fish Demonstrates the Minor Contribution of Coral Mutualists to Their Highly Partitioned, Generalist Diet.” *PeerJ* 3: e1047.

Leray, Matthieu, Joy Y Yang, Christopher P Meyer, Suzanne C Mills, Natalia Agudelo, Vincent Ranwez, Joel T Boehm, and Ryuji J Machida. 2013. “A New Versatile Primer Set Targeting a Short Fragment of the Mitochondrial COI Region for Metabarcoding Metazoan Diversity: Application for Characterizing Coral Reef Fish Gut Contents.” *Frontiers in Zoology* 10 (1): 34.

Mazerolle, Marc J. 2019. “AICcmodavg: Model Selection and Multimodel Inference Based on (Q)AIC(C).” R package version 2.3.3, https://cran.rproject.org/package=AICcmodavg

Schnell, Ida Baerholm, Kristine Bohmann, and M Thomas P Gilbert. 2015. “Tag Jumps Illuminated--Reducing Sequence-to-Sample Misidentifications in Metabarcoding Studies.” *Molecular Ecology Resources* 15 (6): 1289–1303.

Wangensteen, Owen S., and Xavier Turon. 2017. “Metabarcoding Techniques for Assessing Biodiversity of Marine Animal Forests.” In *Marine Animal Forests: The Ecology of Benthic Biodiversity Hotspots*, 445–73. Springer International Publishing.
